# PEPseq Quantifies Transcriptome-Wide Changes in Protein Occupancy and Reveals Selective Translational Repression After Translational Stress

**DOI:** 10.1101/2022.07.28.501819

**Authors:** Jakob Trendel, Etienne Boileau, Marco Jochem, Christoph Dieterich, Jeroen Krijgsveld

## Abstract

Post-transcriptional gene regulation is accomplished by the interplay of the transcriptome with RNA-binding proteins, which occurs in a dynamic manner in response to altered cellular conditions. Recording the combined occupancy of all proteins binding to the transcriptome offers the opportunity to interrogate if a particular treatment leads to any interaction changes, pointing to sites in RNA that undergo post-transcriptional regulation. Here, we establish a method to monitor protein occupancy in a transcriptome-wide fashion by RNA sequencing. To this end, peptide-enhanced pull-down for RNA sequencing (or PEPseq) uses metabolic RNA labelling with 4-thiouridine (4SU) for light-induced protein-RNA crosslinking, and N-hydroxysuccinimide (NHS) chemistry to isolate protein-crosslinked RNA fragments across all long RNA biotypes. We use PEPseq to investigate changes in protein occupancy during the onset of arsenite-induced translational stress in human cells and reveal evidence for ribosome stalling and depletion from stress granules for a distinct set of mRNAs, many coding for ribosomal proteins. We use quantitative proteomics to demonstrate that translation of these mRNAs remains repressed during the initial hours of recovery after arsenite stress. Thus, we present PEPseq as a discovery platform for the unbiased investigation of post-transcriptional regulation.

## Introduction

Protein-RNA interactions organize many cellular processes, ranging from transcription, splicing and translation of protein-coding RNA, to a multitude of regulatory functions organized by small and long non-coding RNA that we are only beginning to understand (1). Protein-RNA interactions are not only vital for the basal cellular operation under homeostatic conditions, but they also allow cells to respond to environmental changes and elicit a commensurate response. In fact, post-transcriptional regulation can effectuate fast changes in gene expression that can often not be realized in time on a transcriptional level. For example, regulation at the level of protein translation occurs in human cells during hypoxia, when non-ribosomal RNA-binding proteins organize the recruitment of the translation machinery to a specific set of messenger RNAs (mRNAs) coding for growth factor receptors and metabolic proteins, ultimately leading to a swift metabolic switch (2, 3). Investigating the binding events occurring for any of these proteins usually requires prior knowledge of their identity (4). For instance, in the above example RBM4 was identified as an interactor of HIF-2α during hypoxia before crosslinking and immunoprecipitation followed by sequencing (CLIPseq) (5), confirmed its binding to a particular motif on the mRNAs that are regulated by the complex between the two proteins (2). Later it was demonstrated that beside RBM4 multiple other RNA-binding proteins influence translation during hypoxia, confirming the notion that translation regulation in response to a stimulus that affects the entire cell involves a superposition of many altered protein-RNA interactions (6–8). Significant efforts have been invested into mapping the transcriptome-wide interaction sites of hundreds of human RNA-binding proteins with CLIPseq as part of the ENCODE project (9). In cases where a biological process is known to involve particular RNA transcripts, this information can be leveraged to identify potential RNA-binding proteins that might bind and affect their function. Yet, for the unbiased investigation of a post-transcriptional event involving unknown proteins and RNAs, it is desirable to map the interaction sites of all RNA-binding proteins across the transcriptome to pinpoint positions where changes in protein occupancy occur during a cellular perturbation. Such data can directly identify RNAs that underlie cellular adaptation, and subsequently be intersected with other sequencing data derived by CLIPseq, Ribo-seq or similar in order to determine what RNA-binding proteins contributed to the changed protein occupancy. Previous methodologies for mapping protein occupancy across the transcriptome use two main approaches – one involving metabolic labelling of the RNA and another one using *in situ* chemical crosslinking. For metabolic labelling of RNA, human cells are cultured in the presence of 4-thiouridine (4SU), a photoactivatable nucleotide that is readily incorporated into nascent RNA (10). Short UV irradiation of intact, live cells chemically activates 4SU-labelled sites in the transcriptome to form covalent crosslinks with interacting proteins in the direct vicinity. RNA extracted from such cells will then lead to characteristic errors during the generation of cDNA for sequencing, introducing T to C transitions at the protein-crosslinked, 4SU-labelled site. Methods such as photoactivatable ribonucleotide-enhanced crosslinking and immunoprecipitation (PAR-CLIP) use these diagnostic transitions in RNA sequencing reads to pinpoint protein-RNA interactions sites with quasi nucleotide resolution (11). Similarly, protein occupancy profiling uses poly-dT enrichment to retrieve protein-crosslinked, polyadenylated RNA from lysates of UV crosslinked cells and after sequencing leverages T-C transitions to map protein interaction sites within these transcripts (12). So far, protein occupancy profiling has been used to compare the static protein interactions with mRNA between cell lines.

Alternatively, chemical crosslinking has been used to map protein-RNA interaction sites. One recently reported method, called ribonucleoprotein networks analyzed by mutational profiling (RNP-MaP), first reacts a bi-reactive probe with lysine residues on proteins in live cells, and subsequently uses UV activation to crosslink them to protein-bound RNA (13). Here, too, diagnostic mutations are introduced into RNA sequencing reads when the reverse transcriptase traverses a protein-RNA crosslink, which can be used to pinpoint protein interaction sites in a transcriptome-wide manner.

We recently reported protein-crosslinked RNA extraction (XRNAX) as a universal method for the purification of photo-crosslinked protein-RNA complexes from UV-irradiated cells (14). We used XRNAX along with MS-based proteomics to establish that the RNA-bound proteome is significantly altered during arsenite-induced translational stress, which similarly to hypoxia leads to impaired translation initiation via EIF2α phosphorylation (15). In this study we aimed to develop a complementary RNA sequencing method to characterize the protein-bound transcriptome and monitor protein occupancy during translational stress. Specifically, we sought to investigate where in the transcriptome the most significant changes in protein-RNA interactions occurred when cells undergo arsenite treatment and how these might relate to adaptive cellular mechanisms. To this end we developed PEPseq (for peptide-enhanced pull-down for RNA sequencing), which uses N-hydroxysuccinimide (NHS) chemistry to exploit the presence of primary amines on peptide-crosslinked RNA to enrich protein-RNA interaction sites and detect them by RNA sequencing. PEPseq employs XRNAX as a starting point for the initial isolation of photo-crosslinked protein-RNA hybrids, which opens the possibility to analyse protein-RNA complexes with a proteomic or transcriptomic read-out from the same sample. Applying PEPseq to human MCF7 cells during the course of arsenite-induced translational arrest we reveal increased protein occupancy across the coding sequences of specific mRNAs, and we use Ribo-seq data to confirm extensive ribosome stalling during translational stress. We find that mRNAs particularly affected by this are strongly excluded from stress granules and encode proteins involved in translation, especially cytosolic ribosomal proteins. Using proteomics, we demonstrate that after arsenite washout and during the recovery from stress, protein production from these mRNAs remains repressed whereas translation of transcripts with strong stress granule localization is prioritized.

## Materials and Methods

### Cell culture and SILAC

All experiments were performed with MCF7 (ATCC, RRID:CVCL_0031) cells maintained in Dulbecco’s Modified Eagle’s Medium (DMEM) supplemented with 10% FBS and Pen-Strep (100 U / ml penicillin, 100 mg / ml streptomycin, Gibco 15140-122) at 37 °C, 5 % CO_2_. DMEM for SILAC (Silantes 280001300) was supplemented with 10 % dialyzed FBS (Gibco 26400-044), 1 mM L-lysine Silantes 211104113, 201204102) and 0.5 mM L-arginine (Silantes 211604102, 201604102) for SILAC intermediate or heavy labels, respectively. Additionally, 1.7 mM light L-proline and 1 x GlutaMAX (Gibco 35050061) were added. Complete SILAC labels were introduced during six passages in the respective SILAC media. Experiments were performed at 70 % confluence and always three days after seeding.

### Peptide-enhanced pull-down for RNA sequencing (PEPseq)

For PEPseq two 24 cm x 24 cm square culture dishes were seeded with one confluent 15 cm dish of MCF7 cells three days prior to arsenite treatment. One day before arsenite treatment 4-thiouridine (biomol, Cay-16373) was added at 100 μM concentration. On the treatment day sodium arsenite (Santa Cruz, sc-301816) was added to a final concentration of 400 μM and cell culture continued normally for 15 or 30 minutes, respectively. The media was then discarded and cells washed twice with ice-cold PBS. The dish was put on ice and UV-irradiated with 0.2 J/cm^2^ in a Vilber Bio-Link mounted with 365 nm UV lamps. Cells were immediately covered in 10 ml PBS and harvested by scraping. Residual cells were collected in another 10 ml PBS and combined in one tube (40 ml total from two dishes) to be spun down at 4 °C with 1000 g for 5 minutes. All PBS was removed and protein-crosslinked RNA extracted by XRNAX as reported before.(Trendel et al., 2019) The entire XRNAX extract was taken up in 1 ml of trypsin digestion buffer (50 mM Tris-Cl pH=7.4, 0.1 % SDS), denatured at 85 °C for 2 minutes and allowed to reach room temperature. Subsequently, 10 μg of trypsin/LysC (Promega, V5073) were added and samples incubated for four hours at 37 °C, 700 rpm shaking. Peptide-crosslinked RNA was then purified with the RNeasy Midi Kit (Quiagen, 75144) according to the manufacturer’s instructions and eluted into 250 μl water. Thirty microliter PBS 10 x concentrate (Calbiochem, 524650) was added, samples brought to 300 μl total volume and heated to 85 °C for 2 minutes before cooling on ice. Samples were then divided in two microTUBEs with AFA fiber (Covaris, 520045) and sonicated on a Covaris S220 (peak power 175, duty factor 50, cycles / burst 200, average power 87.5) for 900 s at 4 °C. Samples were recombined and for the matched input control 30 μl aliquoted and stored at -80 °C before further processing. The remaining sample was brought to a final 400 μl of PBS containing 0.1 % SDS. Again, samples were heated to 85 °C for 2 minutes and allowed to reach room temperature. Per sample 100 μl of NHS-activated magnetic beads (Pierce, 88826) was washed three times with 1 ml and reconstituted in 100 μl PBS containing 0.1 % SDS. Beads were combined with the samples and coupling allowed to occur in a total volume of 500 μl on a rotating wheel overnight. Beads were washed 3 times with 1 ml wash buffer 1 (Tris-Cl 50 mM, 0.1 % SDS) and 3 times with wash buffer 2 (Tris-Cl 50 mM), each time incubating samples for 5 minutes at 55 °C, 700 rpm shaking. In order to introduce the right phosphorylation status on the RNA fragments for sequencing library preparation, beads were treated with 10 μl T4 PNK (Thermo, EK0031) in 90 μl 1 x PNK buffer for 30 minutes at 37 °C, 700 rpm shaking. Beads were washed again twice with 1 ml wash buffer 1 and RNA fragments eluted with 20 μl proteinase K (Thermo, EO0491) in 80 μl proteinase K buffer (Tris-Cl 50 mM, EDTA 10 mM, NaCl 150 mM, SDS 1%) for 30 minutes at 55 °C, 700 rpm shaking. RNA aliquoted for the matched input control was treated with 10 μl PNK in a total volume of 50 μl of 1 x PNK buffer for 30 minutes at 37 °C, 700 rpm shaking. Ten microliters proteinase K were added along with 40 μl 2 x proteinase K buffer and samples digested for 30 minutes at 55 °C, 700 rpm shaking. Both pull-down and input were cleaned up with the RNeasy Mini Kit (Quiagen, 74106) according to the manufacturer’s instructions and eluted into 30 μl water. RNA in the input sample was quantified on a NanoDrop photometer and 1 μg used for sequencing library preparation. The pull-down was used unquantified and the maximum volume applied undiluted for sequencing library preparation. Sequencing library preparation occurred with the NEXTflex Small RNA Sequencing Kit (Perkin Elmer) according to the manufacturer’s instructions using the protocol for long reads and gel-based selection.

### Processing of RNA sequencing and reference data

RNA sequencing reads were pre-processed to make use of the unique molecular identifiers introduced by the NEXTflex Kit, which includes 4 random nucleotides on each side of the read adjacent to the adapter sequence. Therefore, 3’ sequencing adapters (TGGAATTCTCGGGTGCCAAGG) were trimmed with Cutadapt (Martin, 2011) and reads were deduplicated. Subsequently, random 4-mers were removed from the 3’ and 5’ end, and reads shorter than 18 nucleotides discarded. Sequences for ribosomal RNA (NR_003286.4, NR_003285.3, NR_003287.4) were added to the human GENCODEv31 transcriptome (CHR) and annotated as such in the accompanying GTF file. Reads were then mapped to the GENCODEv31 transcriptome (CHR&rRNA) using HISAT2 and applying the forward (F) option, which maps reads only to the actual reference sequence and not its reverse complement. In order to make the differential quantification with PEPseq as straightforward as possible, we applied a two-step mapping strategy that aimed to identify the one transcript that represents all products of a gene in an optimal way (Figure S1C). Therefore, we created a reduced transcriptome to which reads in our entire experiment (pull-down and input) mapped best. We defined two easy criteria all transcript isoforms of a gene were scored by, i) number of mapped reads ii) length. After mapping reads to the entire GENCODEv31 transcriptome, reads were counted (samtools index)(Danecek et al., 2021). Then, for each gene in the GENCODEv31 (CHR&rRNA) annotation we selected the transcript with the most mapped reads, and, in case of a draw, the shortest one. This resulted in our reduced transcriptome (GENCODEv31 transcriptome reduced) used throughout the rest of our analysis. For our transcriptomic analysis we again used HISAT2 and identical settings to map reads again to the reduced transcriptome. From this alignment all following analyses were performed. For our proteomic analysis we translated coding transcripts in the reduced transcriptome (annotated with a CDSs in the GENCODEv31 annotation) to protein and used this as a reference database for the search of our proteomic data.

### Quantitative analysis of PEPseq data

In order to make PEPseq available for a wide audience we used two simple samtools (Danecek et al., 2021) commands to count reads or T-C transitions and DESeq (Love et al., 2014) for differential testing. Bam files with reads mapped against the reduced GENCODEv31 transcriptome were analyzed with samtools to yield read counts per transcript (samtools idxstats) or T-C transitions at a particular position in the transcriptome (samtools mpileup with vcf output). To retrieve counts for the individual mRNA regions bam files were split with samtools (samtools view with bed file containing GENCODEv31 region information) and reads were counted individually in the resulting bam files (samtools idxstats). For the quantification of changes in protein occupancy in the 18S, 28S and 5S rRNA, T-C frequencies at each nucleotide position were applied as one combined input for a DESeq analysis between two time points. An interaction term was used to take T-C frequencies of the pull-down and the input into account during differential testing (∼ experiment + arsenite + experiment:arsenite) and fold changes were corrected with apeglm (Zhu et al., 2019) using the differential effect (experimentpull_down.arsenitetreated). The same model was used to identify differentially occupied transcripts using read counts. Again we used apeglm but in this case tested for the combined effect (arsenite_treated_vs_untreated) as well as the differential effect (experimentpull_down.arsenitetreated). Details about this are discussed in the main text.

### Analysis of published datasets

For our re-analysis of the Ribo-seq data from arsenite-treated HEK293T cells (Ichihara et al., 2021), raw data with the accession number PRJNA729461 corresponding to SRR14510024, SRR14510025 (Ribo-seq), and SRR14510056, SRR14510057 (Ribo-seq, arsenite treatment) were downloaded from the NCBI BioProject database. Reads aligning to a custom bowtie2 v2.3.0 (Langmead and Salzberg, 2012) ribosomal index were discarded. Remaining reads were then aligned in genomic coordinates to the human genome (GRCh38.p13) with STAR v2.5.3a (Dobin et al., 2013) using the following options: ’--quantMode TranscriptomeSAM --alignIntronMin 20 --alignIntronMax 100000 -- outFilterMismatchNmax 1 --outFilterMismatchNoverLmax 0.04 --outFilterType BySJout -- outFilterIntronMotifs RemoveNoncanonicalUnannotated --sjdbOverhang 33 -- seedSearchStartLmaxOverLread 0.5 --winAnchorMultimapNmax 100’. We used evidence from periodic fragment lengths only. Therefore, for each sample individually, fragment lengths and ribosome P-site offsets were determined from a metagene analysis using uniquely mapped reads with our previously developed tool Ribosome profiling with Bayesian predictions (Rp-Bp) (Malone et al., 2017). Fragments with lengths 21 and 28 were extracted from the transcriptome alignment files, and summary statistics calculated with samtools idxstats (Danecek et al., 2021). In order to avoid redundant use of read counts, for each ENSEMBL gene ID, the transcript with the most reads across all samples was selected as the representative isoform. Ratios of translation efficiencies were adapted from the original publication as reported in Table_S7 (Ichihara et al., 2021). To avoid redundancy and give the most conservative representation of translation efficiencies after arsenite stress, only the transcript with the largest ratio for each gene name was used.

For our re-analysis of the stress granule transcriptome from U2OS cells, FPKM values and ratios were adapted from the original publication as reported in Data S1 (Khong et al., 2017). For each ENSEMBL gene id, only the transcript with the highest FPKM value in the total transcriptome was used as the representative isoform. We note here that no filtering led to a very similar outcome.

### Pulsed-SILAC-AHA labeling for translatome analysis during stress recovery

Fifteen centimeter dished were seeded with 1.3 million fully SILAC intermediate-labeled MCF7 cells and cultured for 3 days until 80% confluent. The old media was removed and cells were washed twice with. Cells were accustomed to AHA by incubating them 30 minutes in AHA-SILAC intermediate DMEM (reduced component DMEM (AthenaES 0430), sodium bicarbonate 3.7 g/l, Sodium Pyruvate 1 mM, HEPES 10 mM, GlutaMax 1 x, L-proline 300 mg/l, L-cystine 63 mg/L, L-leucine 105 mg/l, dialyzed FBS 10 %, L-lysine 146 mg/l, L-arginine 84 mg/l, L-azidohomoalanine 18.1 mg/l (Click Chemistry Tools, 1066-1000)). Half of the cultures were treated with 400 μM sodium arsenite for another 30 min, the other half was kept untreated. The old media was removed and cells were washed twice with PBS. The treated cells were then pulsed in DMEM AHA-SILAC-heavy medium, the untreated cells in DMEM AHA-SILAC-light medium for the time points 15, 30, 60, 120 and 180 minutes. At the same time a second replicate with SILAC label swap was produced. The media was discarded and cells were washed with ice-cold PBS. Cells were harvested into 2 × 10 ml ice-cold PBS by scraping and collected by centrifugation. Supernatants were discarded and cell pellets stored at -20 °C until lysis. AHA-labelled proteins were enriched using the Click-it Protein Enrichment kit (Invitrogen C10416) according to the manufacturer’s instructions. Proteins were digested from the beads with 200 μl digestion buffer (tris-Cl 100 mM, acetonitrile 5 %, CaCl_2_ 2 mM) containing 500 ng trypsin/LysC overnight at 37 °C, 1000 rpm shaking in 2 ml tubes. Beads were pelleted by centrifugation and the digest transferred to a new vial. Residual peptides were flushed off the beads with additional 600 μl of water and combined with the digest. Peptides were desalted with an Oasis PRiME HKB μElution Plate and analysed on a QExactive HF HPLC-MS (Thermo Scientific). Separation by HPLC prior to MS occurred on an Easy-nLC1200 system (Thermo Scientific) using an Acclaim PepMap RSCL 2 μM C18, 75 μm x 50 cm column (Thermo Scientific) heated to 45 °C with a MonoSLEEVE column oven (Analytical Sales and Services). Buffer A was 0.1 % formic acid, buffer B was 0.1 % formic acid in 80 % acetonitrile. The gradient used was: 0 minutes 3% B, 0-4 minutes linear gradient to 8 % B, 4-6 minutes linear gradient to 10 % B, 6-74 minutes linear gradient to 32 % B, 74-86 minutes linear gradient to 50 % B, 86-87 minutes linear gradient to 100 % B, 87-94 minutes 100 % B, 94-95 linear gradient to 3 % B, 95-105 minutes 3 % B. The MS detection method was MS1 detection at 120000 resolution, AGC target 3E6, maximal injection time 32 ms and a scan range of 350-1500 DA. MS2 detection occurred with stepped NCE 26 and detection in top20 mode with an isolation window of 2 Da, AGC target 1E5 and maximal injection time of 50 ms.

### MS database search and data analysis

Mass spectrometry data was search with MaxQuant v1.6.0.16 (Cox and Mann, 2008) using the translatome produced from the reduced GENCODEv31 transcriptome (see Processing of RNA sequencing and reference data above) and the default Andromeda list of contaminants. Settings were left at their default values except for the SILAC configurations and activation of the match-between-runs as well as the requantify feature.

For the proteomic data analysis the MaxQuant proteinGroups.txt output was used. Filtering occurred for the columns ‘Potential contaminants’, ‘Reverse’ and ‘Only identified by site’. For the quantification of nascent protein after arsenite stress, unnormalized SILAC heavy / intermediate and light / intermediate ratios were used. Within each time point replicates were combined by inverting one the swapped label and calculating the mean between the ratios. In order to exclude outliers within each time point, filtering occurred for proteins with a standard error of the mean (SEM) larger 30 %.

### Visualization of the 80S ribosome

The cryo-EM structure of the human 80S ribosome (Natchiar et al., 2017) (PDB 6QZP) was accessed in PyMOL (Molecular Graphics System, Version 2.0 Schrödinger, LLC). The peptide exit tunnel was visualized with the coordinates previously reported for this structure (Duc et al., 2019).

### Gene ontology (GO) enrichment analysis

Ranked GO enrichment analysis was performed using the GOrilla web interface (Eden et al., 2009) and ENSEMBL transcript IDs used as input.

### Functional annotation of transcripts and proteins

Information about transcript biotypes was extracted from the GENCODEv31 annotation. ENSEMBL BioMart was accessed via biomaRt and GO annotations used for the functional annotation of proteins.

### Statistical analysis and data visualization

All data handling apart from what is mentioned above was performed in R (4.1.2) with RStudio (1.4.1103) and visualized using the ggplot2 (Wickham, 2016) library. Figures were arranged in Adobe Illustrator (26.0.2). The scheme in Figure 1A was generated with Biorender.

**Figure 1:**
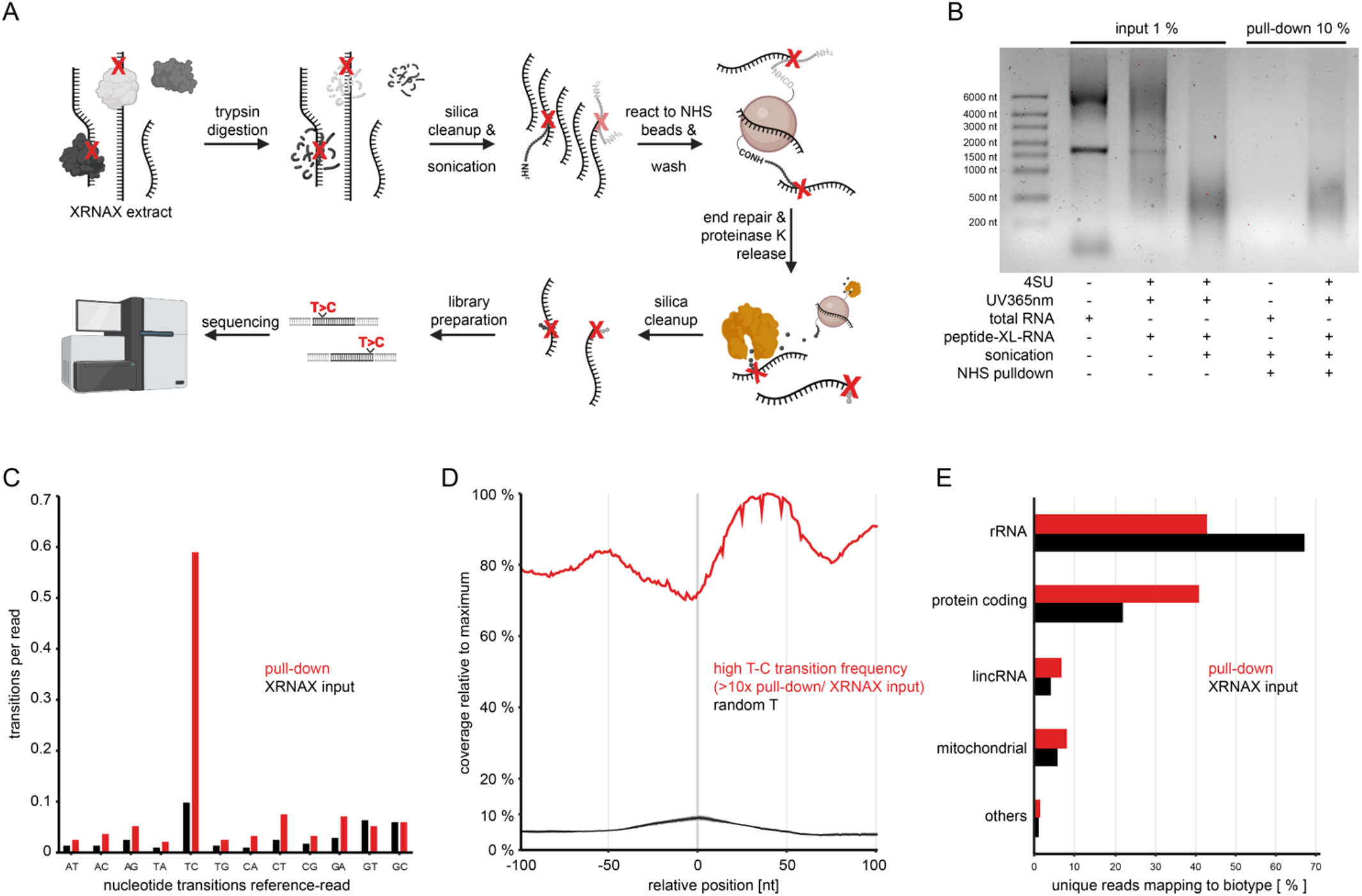
PEPseq captures RNA fragments UV-crosslinked to protein interactors. A) Experimental outline for peptide-enhanced pull-down for RNA sequencing. Cells are 4SU labeled for 24 hours, UV irradiated and protein-crosslinked RNA extracted with XRNAX. Trypsin digestion of the XRNAX input leaves behind peptides crosslinked to the RNA, whereas non-crosslinked peptides are removed using conventional silica columns. Ultrasonicated RNA fragments are applied for the pull-down and also sequenced as matched input control. For the pull-down primary amines are reacted to NHS-activated beads thereby covalently capturing fragments carrying crosslinking sites. Elution occurs with proteinase K, cleaving captured fragments off the beads while avoiding harsh elutions that release non-covalently captured background fragments. B) Agarose gel electrophoresis comparing the NHS-mediated pull-down of protein-crosslinked RNA and non-crosslinked, protein-free RNA. Protein-free RNA (total RNA) was extracted from untreated MCF7 cells using standard silica spin columns with additional proteinase K digestion. The preparation of peptide-crosslinked RNA (peptide-XL-RNA) and the NHS-mediated pull-down occurred as described in A. C) Bar graph comparing nucleotide transition frequencies in unique sequencing reads from peptide-enriched fragments (pull-down) and the matched inputs they were extracted from (XRNAX input). D) Linegraph illustrating the combined read coverage around positions in the transcriptome with T-C transition frequencies ten times higher in the pull-down than in the input (high T-C transition frequency, red line). As a control the combined read coverage around the identical number of random Ts is indicated (black line indicating the mean, grey shading the standard deviation of 200 randomizations). See also Figure S1B. E) Bar graph comparing read counts within GENCODE RNA biotypes.

## Results

### Utilizing NHS-Chemistry to Capture Peptide-RNA Hybrids for RNA Sequencing

We previously developed XRNAX to purify UV-crosslinked protein-RNA complexes from human cells, allowing their characterization both from a proteomic and transcriptomic perspective. Sequencing of protein-crosslinked RNA that was extracted via XRNAX from 4SU-treated MCF7 cells resulted in an increased T-C transition frequency (14), yet, not to the degree that permitted mapping of protein occupancy in a meaningful way (data not shown). This indicated that an additional enrichment step was necessary to selectively extract RNA fragments carrying UV crosslinking sites, and to eliminate non-crosslinked RNA. Protein-crosslinked RNA that is extracted with XRNAX, digested with proteinase K and cleaned up with silica spin columns, is mostly protein-free with sporadic modifications by crosslinked peptide residues. We exploited the presence of a primary amine in such peptide remnants, which is absent in non-crosslinked RNA, by reacting NHS-biotin with RNA prepared via XRNAX, for subsequent enrichment of peptide-RNA hybrids on streptavidin beads. RNA sequencing confirmed that diagnostic T-C transitions were strongly increased in NHS-biotin-enriched RNA fragments compared to the XRNAX input, however, only in a minority of reads (<30 %, data not shown). Because the streptavidin-biotin interaction is very stable, biotinylated peptide-RNA hybrids needed to be released by boiling the beads in formamide. We hypothesized that this also released non-crosslinked RNA fragments covering the beads, which normal washing had not been able to remove effectively. This hypothesis was supported by the failure of various washing buffer formulations to reduce the background. Hence, instead of using proteinase K, which cleaves after any amino acid in a peptide sequence, we switched to trypsin for the initial digestion step, which only cuts after lysine or arginine and therefore leaves longer peptides attached to a UV-crosslinked RNA fragment. Tryptic peptides carry primary amines only at their termini so that the peptide sequence in between remained as a spacer that was accessible to other peptidases. This meant we could omit biotinylation, but instead covalently couple peptide-bearing RNA fragments immediately to NHS-activated beads. Subsequently, we used proteinase K at moderate conditions to digest the remaining amino acid sequence and liberate the RNA fragment from the bead (Figure 1A). This enzymatic elution strongly decreased the background so that virtually no RNA could be enriched from non-crosslinked, protein-free RNA, whereas efficient enrichment occurred from protein-crosslinked and trypsin-digested RNA (Figure 1B). Moreover, RNA sequencing demonstrated that this strategy greatly increased reads with T-C transitions, which occurred now in more than 60 % of reads (Figure 1C). Notably, most reads in the pull-down without a T-C transition carried some other nucleotide transition, implying that a peptide-RNA crosslink had interfered with reverse transcription somehow (5). In turn this indicated that our new elution strategy had greatly reduced the background of non-crosslinked RNA fragments and allowed us to sequence fragments strongly enriched in protein-RNA interaction sites. We named this new method peptide-enhanced pull-down for RNA sequencing (PEPseq), which we aimed to apply for the transcriptome-wide mapping of protein-RNA interaction sites. T-C transitions occurred most frequently towards the 3’ and 5’ ends of reads (Figure S1A). This may be caused by the sterical properties of RNA fragments, which might be more reactive when carrying a crosslinking site towards their ends. Increased frequency of T-C transitions (T-C count normalized by coverage at position) in the pull-down compared to the input was also evident at individual transcriptome positions (Figure S1B), although some positions had a higher background of T-C transition frequencies than others. Since our biological interpretation of an increased T-C transition frequency was protein binding, this background needed to be considered during our following analysis. Conceptually, PEPseq offers two read-outs for protein-RNA interactions, namely the count of diagnostic T-C transitions in reads covering a certain position in the transcriptome, and the read-count itself. As calculating meaningful T-C transition frequencies requires robust coverage of a transcript by many reads per position, simply counting reads might include transcripts with low coverage in the analysis, for which no reliable T-C transition frequency would normally be calculated. To validate pull-down reads in PEPseq as an indicator for protein occupancy, we selected transcriptome sites where the T-C transition frequency was at least ten times higher in the pull-down than in the input, i.e. high-confidence protein-RNA interaction sites (Figure S1B). Figure 1D shows that the read coverage around these sites was on average more than one order of magnitude higher than at random control sites, strongly indicating that pull-down reads in PEPseq libraries were a proxy for T-C transitions in marking protein occupancy. Earlier studies revealed that a static view on protein occupancy across mRNA gives little insight into post-transcriptional processes (12, 16). Since we aimed to design PEPseq for the quantification of changes in protein occupancy between conditions, we therefore introduced three design paradigms to harmonize our experimental setup with the subsequent computational analysis and differential testing (Figure S1C). First, since the sequence complexity was greatly reduced in pull-down samples we used unique molecular identifiers (UMIs) to identify true replicate reads and exclude PCR replicates. Second, we sequenced the input from which the pull-down occurred in the exact same way as the pull-down, thus increasing the power of subsequent statistical analyses. Lastly, we limited our reference transcriptome to one transcript per gene, and extended this principle in our proteomics follow-up experiments to one gene – one transcript – one protein. This greatly simplified and strengthened our quantification because no statistical power was lost to isoform information, which we did not intend to use anyway. We note here that because of this one gene – one transcript – one protein approach, we use for clarity ‘gene’ in the following as an umbrella term to refer to any of them.

In summary, we utilize XRNAX and NHS click-chemistry to covalently enrich peptide-RNA hybrids for RNA sequencing in PEPseq – a dedicated method for the quantification of changes in protein occupancy across the transcriptome.

### Translation Arrest Induces Distinct Changes in Protein Occupancy Within mRNA Regions

To benchmark PEPseq we applied it to a system where robust changes in protein-RNA interactions were expected. Since mRNAs are continuously traversed by translating ribosomes we anticipated that inhibiting translation should lead to a strong change in their interaction with ribosomal proteins. Hence, we inhibited translation in MCF7 cells to investigate i) if PEPseq was able to detect changes in protein-RNA interactions across mRNA, ii) if this change differed between coding and non-coding regions in mRNAs, and iii) if alterations in protein-RNA interactions could be observed for non-coding RNAs. In order to follow the change in protein occupancy over time we chose arsenite as a translation inhibitor, which, unlike antibiotic inhibitors of the ribosome, does not lead to immediate translational arrest but progressive translational shutdown within 30 minutes of treatment (14, 17). We created PEPseq sequencing libraries comprising pull-downs and matched inputs from biological duplicates of untreated MCF7 cells, and cells treated with arsenite for 15 or 30 minutes. PEPseq reads mapped to 23,766 unique genes, 67% of which were represented in the input where reads mapped to 35,097 unique genes. Figure 1E shows that PEPseq libraries had a much smaller proportion of reads mapping to ribosomal RNA (rRNA), while they contained a higher proportion of protein-coding and long intergenic non-coding RNA (lincRNA). This effect has been previously observed in other applications that apply 4SU labelling for enrichment of RNA, where it was proposed that ribosomes and rRNA are turned over more slowly than other RNA biotypes and, thus, accumulate a much weaker 4SU label (18). We turned our attention towards mRNA and combined the T-C signal of all detected coding transcripts to monitor common trends across their three functionally defined regions, the 3’ untranslated region (3’ UTR), the protein coding sequence (CDS) and the 5’ untranslated region (5’ UTR). Because longer transcripts have a higher chance of accumulating T-C transitions we normalized the absolute T-C count to the region length of each individual transcript. Indeed, Figure 2A shows that cumulative T-C counts distributed very reproducibly between replicates within each of these regions and showed a distinct change in the 30 minutes time point. In unperturbed cells normalized T-C counts were very similar between the CDS and the 3’ UTR, yet, remarkably lower in the 5’ UTR. Interestingly, we observed that an additional difference emerged between the CDS and the 3’ UTR in cells treated with arsenite for 30 minutes. This clearly recapitulated the three functional mRNA regions, which could now be readily distinguished by their distinct T-C counts. As expected, a similar observation was made when employing read coverage instead of T-C counts (Figure S2A). To better resolve the change over time we normalized cumulative T-C transitions in the pull-down to the input and compared later time points to the untreated sample (Figure 2B & Figure S2B). For cells treated with arsenite for 15 minutes, this indicated that the protein occupancy was overall slightly lower across all mRNA regions compared to untreated cells. For cells treated for 30 minutes, protein occupancy decreased even further, however, only in the 5’ and 3’ UTR and not the CDS. The same analysis using read coverage instead of T-C transitions as read-out for protein occupancy revealed the same effect, even suggesting that the CDS slightly gained protein occupancy 30 minutes into arsenite stress (Figure 2C & Figure S2C).

**Figure 2:**
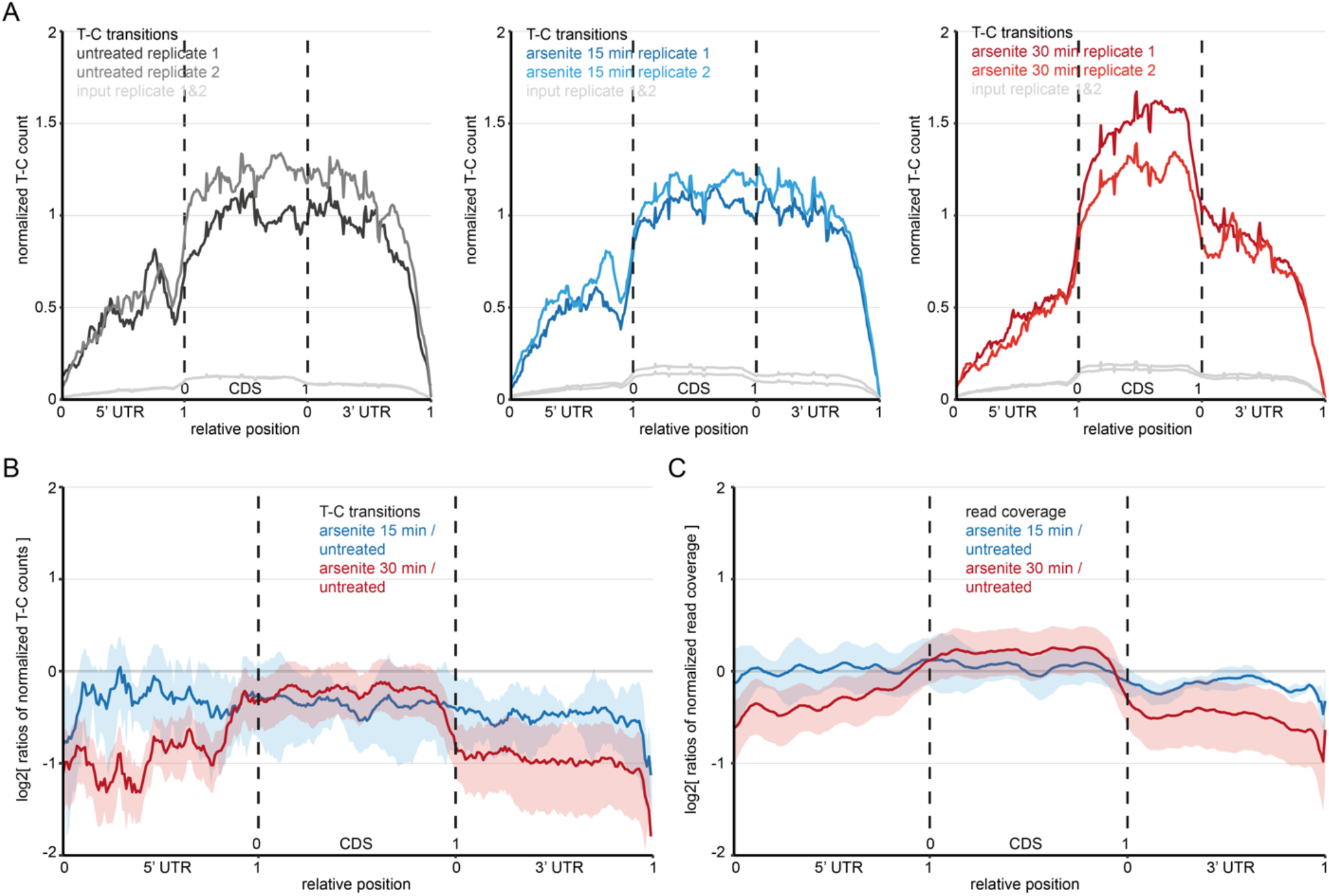
Changes in protein occupancy across mRNA upon arsenite-induced translational arrest. A) Metagene plot summarizing T-C transitions in the 5’ untranslated region (5’UTR), the coding region (CDS) and the 3’ untranslated region (3’UTR) of all detected mRNAs in untreated (left), 15 minutes (middle) and 30 minutes (right) arsenite treated MCF7 cells. The T-C count was normalized to the length of each region for each particular transcript and to the number of reads mapping to mRNA in the individual sample. B) Metagene plot illustrating the change in T-C transitions upon arsenite stress across all detected mRNAs. Ratios of ratios comparing the indicated time point to the untreated control, and normalizing the pull-down to the XRNAX input. Thick lines indicate ratio means, shaded areas one composite standard deviation. See also Figure S2B. C) Same as in B but read coverage is used as proxy for protein occupancy instead of T-C transitions. See also Figure S2C.

These experiments confirmed that PEPseq was able to capture changes in protein-RNA interactions, and locate these changes in the transcriptome in a biologically meaningful way.

### Increased Protein-Occupancy in the Peptide Exit Tunnel Points to Ribosome Stalling

The observed increase in protein occupancy across the CDS during the last half of a 30-minute arsenite treatment (Figure 2) was surprising, as translational arrest should result in fewer ribosomes traversing the CDS, which would intuitively imply reduced protein occupancy. It has been reported that arsenite treatment leads to ribosome stalling on some mRNAs (19), hence, we speculated that translating (i.e. moving) ribosomes might be less susceptible to UV crosslinking than 80S complexes stalling on mRNA. Conversely, stalling ribosomes and the repair machinery they attract might explain increased UV-crosslinked protein in the CDS and increased PEPseq signal in the CDS upon arsenite stress. Using cryo-electron microscopy we recently reported that 80S ribosomes from arsenite-treated MCF7 cells adopt almost exclusively a post-translocation conformation, indicating that their normal ratcheting motion becomes impaired, potentially leading to stalling of ribosomes on mRNA (20). In yeast and human cells it was demonstrated that in ribosome profiling (Ribo-seq) the length of a ribosome footprint informs about the conformation of the ribosome during its ratcheting motion on mRNA (21). To find further evidence that arsenite stalled ribosomes in the CDS, we therefore first turned to a published Ribo-seq dataset investigating arsenite-induced translational arrest in human HEK293 cells (22). Indeed, we observed a strong 2.5 fold shift from 21 nucleotide reads (pre-translocation) to 28 nucleotide reads (post-translocation) upon arsenite treatment in the published Ribo-seq data (two-sided Kolmogorov-Smirnov test p<2E-16, Figure S3A). This aligned with our cryo-EM findings (20), and overall suggested that arsenite stress led to an accumulation of 80S ribosomes trapped in the post-translocation conformation across the CDS. Next, we turned our attention to the ribosomes themselves and asked if there were any indications in our PEPseq data for changes in protein-RNA interactions towards ribosomal RNA (rRNA). Both the pull-down and the input PEPseq libraries had high coverage for ribosomal transcripts, and the ratio of T-C transition frequencies at individual sites of the rRNA (normalizing the pull-down to the input) was remarkably reproducible between replicates (R^2^=0.83, Figure S3B). We observed similar distributions of T-C transition frequencies between the time points, supporting the intuition that the majority of protein-RNA interactions in a stable complex like the ribosome remain constant (Figure S3C). Consequently, we looked for differences in T-C transitions on the nucleotide level in order to identify specific sites within the rRNA where significant changes occurred between the time points. Therefore, we employed the commonly used DESeq2 tool for differential testing (23) and used an interaction term in our model (T-C count ∼ treatment + experiment + treatment:experiment) in order to make use of our matched T-C transition frequencies for both pull-down and input. This revealed one uridine at position 4558 with very significantly changed protein occupancy on the 28S rRNA (Figure 3A, Table S1). A cross-reference with a high-resolution cryo-electron microscopy (cryo-EM) structure of the human ribosome identified this uridine as part of the wall of the peptide exit tunnel (tunnel wall, Figure 3B&C) (24, 25). The site strongly increased its protein occupancy 15 minutes into the experiment (9.3 fold increase over untreated, adj. p=1.4E-24) and remained strongly occupied until completion of the 30 minutes (6.6 fold increase, adj. p=4.4E-5), suggesting that the resting, nascent peptide became especially susceptible to UV crosslinking with the one particular uridine residue in the wall of the peptide exit tunnel (Figure 3C). We also observed clusters of increased protein occupancy at several sites within two stretches of the rRNA (flexible loops, Figure 3A), which were unresolved in the cryo-EM structure. Figure 3B illustrates the 5’ and 3’ regions of these sections forming stumps that protrude from the large subunit into unresolved extensions. While at the moment we cannot say what proteins caused these interactions, they indicated potential regulatory events occurring on the surface of the ribosomal complex.

**Figure 3:**
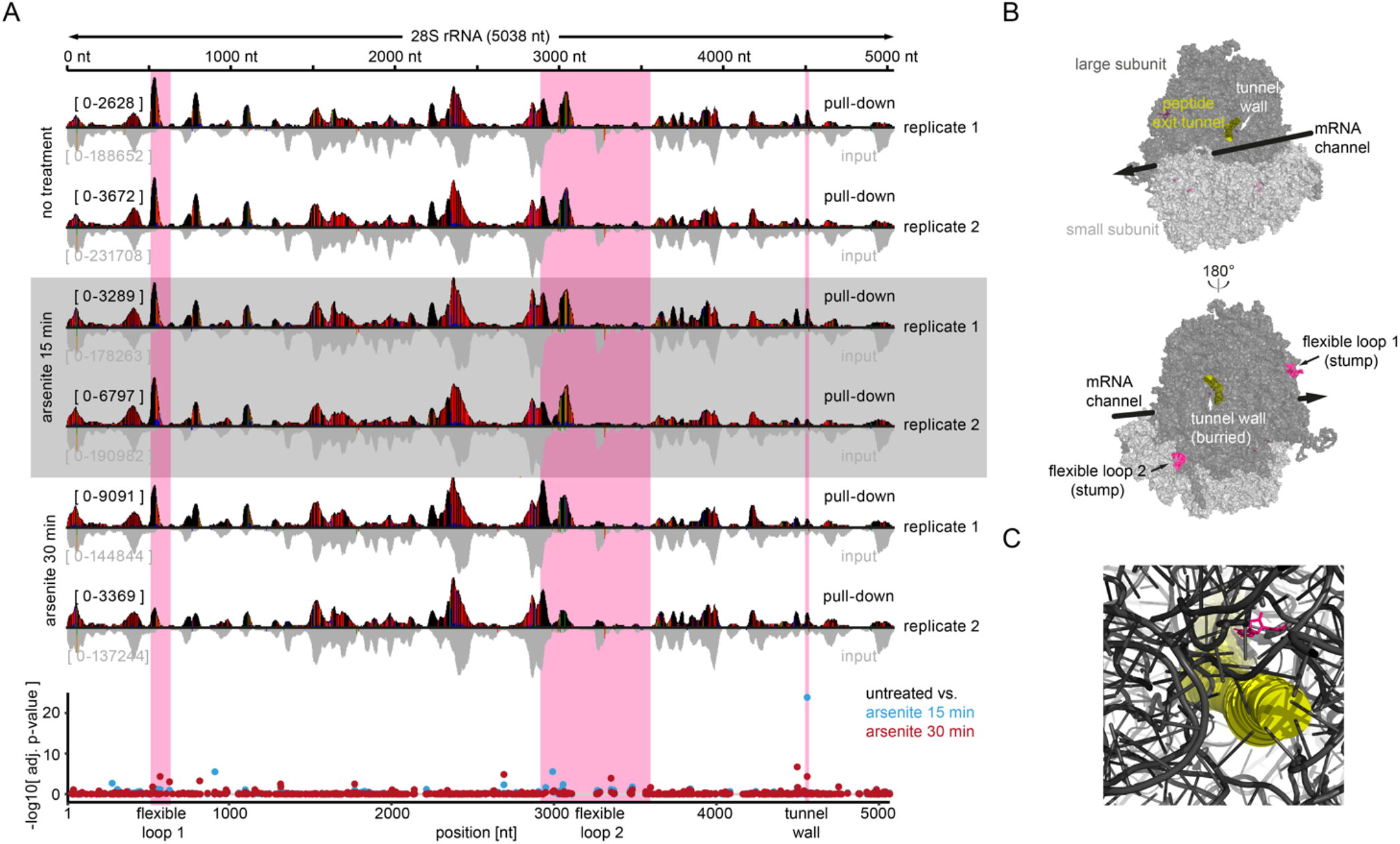
Dynamics of protein interactions with ribosomal RNA upon translational arrest. A) Gene browser view of the 28S rRNA for each replicate pair of pull-down (top, black) and input (bottom, grey). Sites with a T-C conversion frequency higher 5 % are colored in red and blue. Below: Scatter plot indicating significance level of changes in T-C transition frequencies detected by DEseq. Testing occurred between untreated cells and cells treated for 15 minutes (blue) or 30 minutes (red) with arsenite. Magenta shaded areas are highlighted in B. B) Cryo-electron microscopy structure of the human 80S ribosome visualizing differential protein-RNA interactions upon arsenite treatment. The large subunit is shown in dark grey, small subunit in light grey. For orientation the big black arrow indicates the direction of travel for the mRNA. The peptide exit tunnel is filled with yellow cylinders. Regions with significantly changed protein occupancy highlighted in A are shown in magenta. C) Cryo-EM structure of the human ribosome detailing the peptide exit tunnel as shown in the top panel of B. The uridine at the tunnel wall indicated in B is highlighted in magenta stick representation. For visibility all other features are displayed in cartoon representation.

In summary, this analysis showed that T-C transition frequencies in PEPseq data could be leveraged to quantify protein-RNA interactions within abundant transcripts at *quasi* nucleotide resolution, recapitulating the stalling of 80S ribosomes during arsenite-induced translational arrest.

### A Specific Set of mRNAs Becomes Occupied in their CDS During Stress

In order to better understand if certain mRNAs increased their protein occupancy more than others, we next took our analysis towards individual transcripts. Therefore, again DESeq2 was applied to compare the count of reads mapping to individual genes between untreated or arsenite-treated cells by integrating the input-controlled pull-down experiment into one analysis using a model with interaction terms (read count ∼ treatment + experiment + treatment:experiment). The model allowed us to identify two kinds of events, i.e. genes significantly changing in the same direction in both the pull-down and input (combined effect, Figure 4A & S4A, Table S2) or genes significantly changing in the pull-down relative to the input (differential effect, Figure 4B & S4B, Table S2).

**Figure 4:**
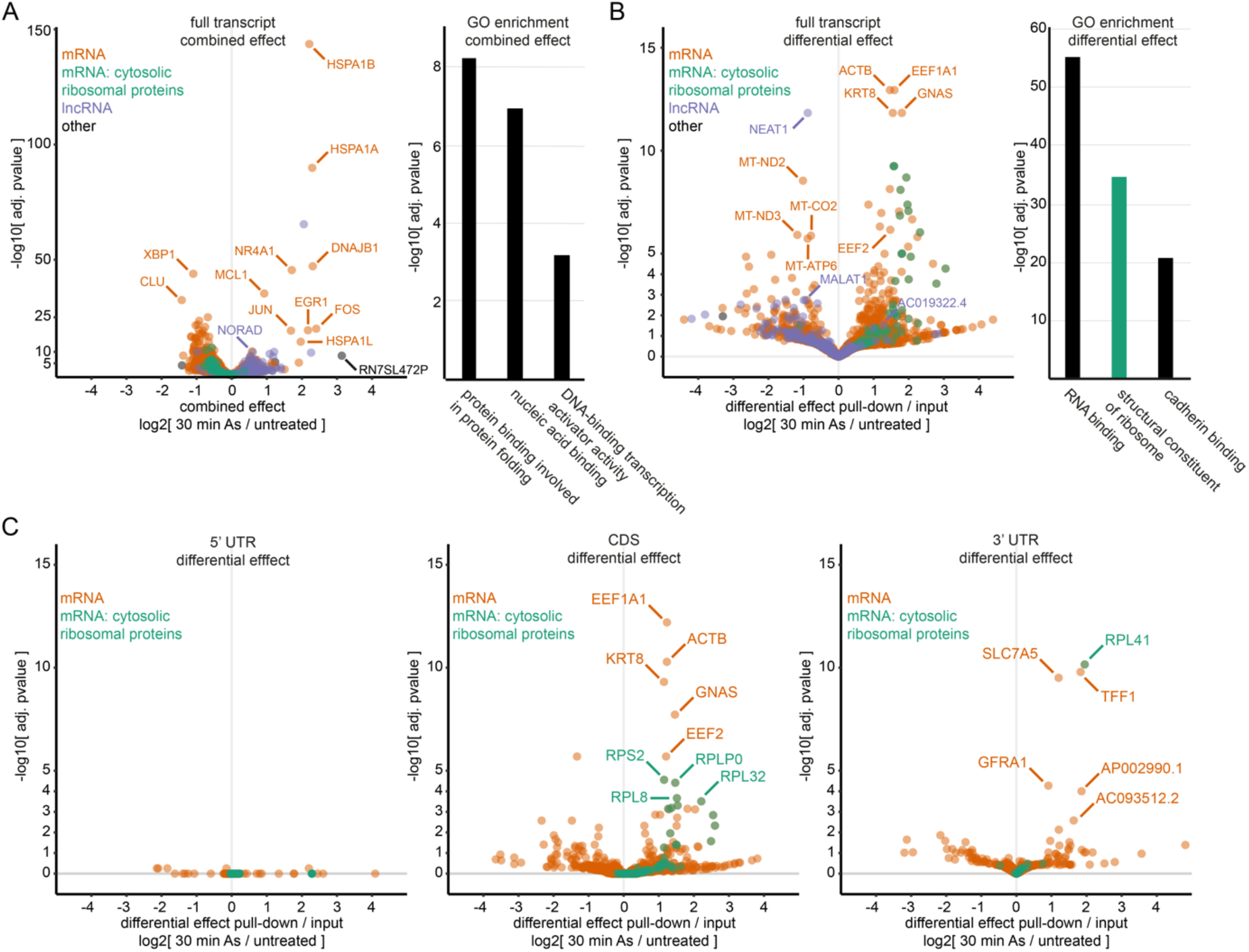
Arsenite stress leads to distinctive changes in protein occupancy across specific transcripts. A) Left: Volcano plot illustrating DEseq results for the combined effect between pull-down and XRNAX input upon 30 minutes arsenite stress. Each point represents one transcript/ gene. Right: Ranked GO enrichment on protein coding transcripts in the DEseq analysis sorted by their foldchange. Shown are the top 3 non-redundant GO terms. B) Same as in A but for the differential effect detected by DESeq between pull-down and XRNAX input. C) Same as in B but for functional regions of protein coding transcripts. DEseq was applied to test for the differential effect within the 5’ UTR (left), CDS (middle) and 3’ UTR (right) using the differential effect between pull-down and XRNAX input. See Figure S4D for exemplary genome browser views.

Genes showing a strong combined effect after 30 minutes of arsenite stress were especially coding for proteins involved in the heat-shock or misfolded protein response. Beside proteins from the actual heat-shock family (HSPA1B, HSPA1A, DNAJB1, HSPA1L), this encompassed the stress-related transcription factors NR4A1, EGR1 and JUN/FOS, which are known ‘immediate-early’ regulators for cell survival (26– 28). Ranked gene ontology enrichment on the list of genes sorted by the combined effect confirmed that the transcriptional profile after 30 minutes arsenite stress was dominated by the misfolded protein response and transcription regulation (Figure 4A). The effect was much less developed at the 15-minute time point, yet, similar tendencies became evident, highlighting an early induction of EGR1 (Figure S4A). As a general trend we observed that the expression of protein-coding genes was slightly reduced, whereas the expression of long non-coding RNAs (lncRNAs) was increased (Figure 4A). Overall, these observations indicated that the combined effect captured the transcriptional response of the cell that progressively increased the production of RNA coding for proteins managing the proteotoxic effects of arsenite.

Next, we examined the differential effect that identified genes where read counts significantly deviated in the pull-down between time points when normalized to the input (Figure 4B & S4B). Since reads in the pull-down mapped protein-RNA interactions (Figure 1D), this implied the differential effect offered an input-controlled way to monitor genes changing their protein occupancy over time. Strikingly, we identified numerous genes coding for proteins involved in translation with significantly increased protein occupancy 30 minutes into arsenite stress (Figure 4B). In fact, gene ontology enrichment on the protein coding genes in the analysis sorted by the 30-minute differential effect returned ‘structural constituent of ribosome’ as the second top ranking term (adj. p= 2.3E-35) only surpassed by ‘RNA-binding’ (adj. p= 8.0E-56). This implied that differentially occupied genes were functionally related, coding for translation-associated proteins and especially cytosolic ribosomal proteins. Interestingly, no significant changes in protein occupancy were detected at the 15 minutes time point (Figure S4B). In order to better understand the altered protein-RNA interaction sites, we repeated the differential DESeq analysis for the individual functional regions of mRNA (Figure 4C). This clearly indicated that the increase occurred in the CDS, which was confirmed by manual inspection of the most strongly affected mRNAs in the genome browser (Figure S4D). In regard to our previous observations on the combination of all mRNA (Figure 2), these additional findings now implied that the increase of protein occupancy across the CDS occurred on specific mRNA species with a common biological function. We used the 30 minutes time point to select sets of genes with significantly (adj. p<0.05) increased protein occupancy (iPO mRNA) or decreased protein occupancy (dPO mRNA) in their CDS for our follow-up analysis.

Collectively, we demonstrate that PEPseq data carries different layers of transcriptomic information, revealing an arsenite-induced transcriptional response geared towards the production of stress-related proteins, as well as potential ribosome stalling on a specific set of mRNAs coding for translation-associated proteins.

### mRNAs with Increased Protein Occupancy in the CDS Are Depleted from Stress Granules

In order to validate the observations we had made for protein coding transcripts, where certain groups of mRNA increased their protein occupancy during arsenite stress, we cross-referenced them with published data from different cell lines. Therefore, we first asked if stalling ribosomes might be the cause for this increased occupancy and re-analyzed Ribo-seq data from arsenite-treated HEK293 cells (22). In this dataset, translation efficiencies of iPO mRNAs decreased strongly upon arsenite (p=2.7E-7, two-sided Kolmogorov-Smirnoff test, Figure S5A). Moreover, ribosome footprints indicative for the post-translocation conformation increased on average twice as much for iPO mRNA during stress compared to all other transcripts (p=2.0E-6, Figure S5B). Both observations confirmed that ribosome stalling was particularly strong on transcripts discovered by PEPseq. Because iPO mRNA comprised many mRNAs coding for ribosomal proteins, we also tested this group in isolation and observed an even stronger effect in both cases (p<8.4E-10, Figure S5B).

Arsenite is commonly used to induce stress granules – large protein-RNA aggregates that form in the cytosol upon translational arrest and whose cellular functions are still poorly understood (29). It has been reported that stalled 80S complexes prevent the recruitment of individual transcripts into stress granules (19). We were interested to see if that was also the case for iPO mRNAs, and turned to a dataset by Khong *et al*., who sequenced the stress granule transcriptome using an immunoprecipitation strategy towards the stress granule marker G3BP1 in arsenite-treated U2OS cells) (30). By comparing read counts from the RNA sequencing of immunoprecipitated stress granules to the total RNA of the same cells (Figure S5C), the authors devised a fold change (in the following referred to as SG score by Khong *et al*.) to rank transcripts according to the degree they became sequestered into stress granules (high SG score) or remained dissolved in the cytosol (low SG score). When applying the SG score by Khong *et al*. to our data, we observed that iPO mRNAs preferentially localized outside of stress granules (two-sided Kolmogorov-Smirnov test, p=1.9E-7), whereas dPOs did not (p=0.39, Figure S5E). Again, mRNAs coding for cytosolic ribosomal proteins in isolation were also strongly excluded from stress granules (p<2.0E-16). Importantly, this was independent of their absolute expression levels in the total transcriptome, spanning more than three orders of magnitude (Figure S5D). Accordingly, ranked GO enrichment analysis on the list of protein-coding genes by Khong *et al*., sorted from most depleted to most included in stress granules, returned ‘structural constituent of ribosome’ as top hit (adj. p=4.3E-59, Figure S5F). The inverted sorting returned ‘DNA binding’ (adj. p=4.0E-15) as top hit and other terms involving regulatory processes such as transcription regulation, GTPase or kinase signalling. This implied that different sets of mRNAs coding for proteins with a shared molecular function were sorted into (or depleted from) a common location upon translational arrest. Supposing that mRNAs in stress granules are stored so they can be subject to particular translation regulation during or after stress (29), this sorting suggested the preparation of a biological program.

This analysis confirmed in a different cell line that iPO mRNA collect stalled ribosomes under arsenite stress and suggested that this might lead to their exclusion from stress granules.

### Protein Production from mRNAs with Increased Protein Occupancy is Repressed After Stress

We hypothesized that stalled ribosomes in the CDS would probably not be an obstacle during arsenite stress because translation would be arrested at the point of initiation (31, 32) and, consequently, no more ribosomes would event try to traverse the mRNA. However, stalled ribosomes would become an obstacle when arsenite stress was overcome, translational arrest was resolved, and mRNAs were translated again. To investigate if this might influence protein expression from specific mRNAs after stress, we designed a proteomic experiment which selectively monitored the reuptake of translation during the recovery from arsenite-induced translational arrest (Figure 5A). Specifically, MCF7 cells carrying an intermediate SILAC label were switched to media containing azidohomoalanine (AHA) instead of methionine for 30 minutes in order to generate an equal amount of SILAC-intermediate-labelled proteins to quantify against. Subsequently, cells were left untreated or exposed to arsenite for another 30 minutes, washed, and switched to AHA-containing media with a heavy SILAC label (for arsenite-treated cells) or light SILAC label (for untreated cells). During a subsequent chase, six time points between 15-240 minutes were collected, and untreated or arsenite-treated cells of each time point were combined to quantify their nascent proteome during this period. Therefore, AHA-containing proteins were enriched by click-chemistry, proteolytically digested, and analysed by mass spectrometry. Depending on the labelling time and treatment this quantified between 1607-3299 proteins at each time point after filtering (REM<30 %, Figure S5G). Ratios between the SILAC intermediate label and the heavy or light label were used to assess which proteins were produced immediately after arsenite stress compared to unperturbed controls (Figure 5B, Table S3). As expected, we observed that during the initial phase after arsenite removal, protein production was weak. Two hours of recovery marked an inflection point where most proteins returned to a production rate similar to the one observed in untreated control cells. Strikingly, protein production from iPO mRNA was reduced compared to the overall protein production during the early time points of recovery and especially 30 minutes after arsenite stress (Figure 5C, two-sided Kolmogorov-Smirnov test with Bonferroni-Holm correction, p=3.4E-4). Because iPO mRNA preferably located outside of stress granules (Figure S5D & Figure S5E), we next asked if in general mRNAs inside or outside of stress granules were translated differently during the recovery from translation arrest. Indeed, at time points immediately after arsenite treatment we found a weak positive correlation between the SG score by Khong *et al*. and the relative protein production (Spearman correlation ρ≤0.28 after 15 minutes, Figure S5H). In contrast, for untreated cells we observed a weak anticorrelation between protein production and stress granule localization throughout all time points. To substantiate this further, we selected the 100 proteins within each time point whose cognate mRNA had the highest or lowest SG score according to Khong *et al*., and compared their relative protein production. This revealed highly significant differences between the two groups (Figure 5D). Especially during the early time points until two hours after arsenite stress, proteins were much stronger produced from mRNA with strong stress granule localization (Wilcoxon ranksum test with Bonferroni-Holm correction, p<1.2E-7 for all time points before 120 minutes). In untreated control cells, we observed the inverse trend, so that transcripts outside of stress granules were stronger translated throughout all time points (p<1.5E-5 for all time points after 30 minutes). In arsenite-treated cells, two hours marked an inflection point where mRNAs with reportedly weak stress granule localization picked up protein production and became significantly more translated than mRNAs with strong stress granule localization (p<3.2E-7 for all time points after 120 minutes).

**Figure 5:**
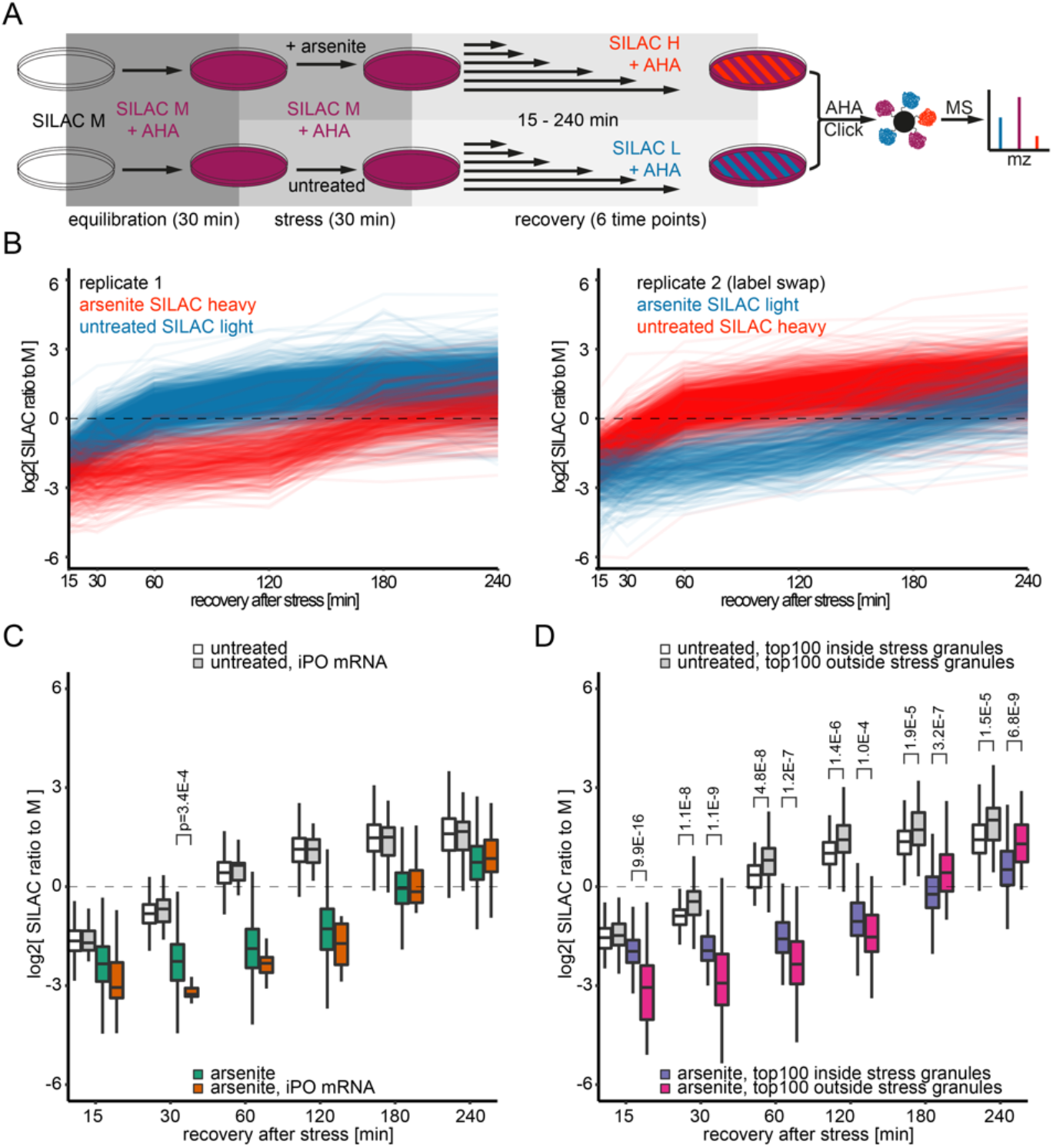
Protein production during the recovery from arsenite-induced translational arrest. A) Experimental scheme for the quantification of nascent protein during recovery from arsenite stress. For details see text. B) Timeline displaying the protein produced in MCF7 cells after 30 minutes of arsenite stress. Each line represents one protein. Only proteins quantified across all time points with a relative standard error of the mean (REM) smaller 30 % are displayed. C) Boxplot comparing the protein production after stress for all detected proteins (arsenite, untreated) to protein produced from iPO mRNAs. Proteins were filtered for REM<30% within each time point and treatment. Testing occurred with a two-sided Kolmogorov-Smirnov test and Bonferroni-Holm correction. D) Boxplots displaying the protein production after stress from transcripts inside or outside of stress granules, respectively. Within each treatment and time point the 100 proteins with the highest/lowest stress granule score according to Khong *et al*. are compared. Filtering occurred as in D, testing with a Wilcoxon ranksum test and Bonferroni-Holm correction.

Next, we asked what sequence features in the iPO mRNAs might lead to ribosome stalling or exclusion from stress granules. Almost all mRNAs coding for cytosolic ribosomal proteins carry a 5’ terminal oligopyrimidine (5’ TOP) motif, which is known to be involved in translational control via interactions with RNA-binding proteins such as LARP1 (33). However, hundreds of other mRNAs strongly excluded from stress granules clearly do not carry the 5’ TOP, so that the motif is not predictive for stress granule depletion. Moreover, it is hard to imagine how a motif in the 5’ UTR would lead to stalling of ribosomes in the CDS and we did not find any other motifs enriched in iPO mRNA. Alternatively, certain sequence features in the CDS are more prone to ribosome stalling than others. In yeast, for example, it was shown that oxidation of G to 8-oxo-G results in very effective ribosome stalling during oxidative stress (34). The occurrence of purine bases in the wobble position (GC3 content) was remarkably high in iPO mRNA (Figure S5I), raising the possibility that the sequence composition of the CDS could be a factor for ribosome stalling. It will require detailed follow-up studies to investigate if this is the case, and whether high GC3 content may regulate SG localisation under translational stress.

Overall, these data imply that mRNA excluded from stress granules during translational arrest, such as iPO mRNA, are translationally repressed during the initial stress recovery, while the opposite is true for mRNA with strong stress granule localization. These genes appear to establish a program to reorganize the cell after stress, representing a previously unrecognized modality of post-transcriptional gene regulation.

### Arsenite-Induced Changes in Protein-RNA Interactions Across Non-Coding RNA

Apart from the observations we made for protein coding transcripts we found profound effects of arsenite on the non-coding transcriptome and its protein occupancy. Among these was the long non-coding RNA NORAD (combined effect adj. p=3.06E-07, Figure 4A), which has been reported to form cytosolic condensates with the PUM family of proteins upon arsenite stress (35). Indeed, we could identify in particular one region in NORAD with an increase in protein occupancy overlapping with published PUM1 and PUM2 CLIPseq data from K562 cells (Figure S6A) (9). When normalized to their total expression in the input most lncRNAs retained or decreased their protein interactions upon arsenite stress (differential effect, Figure 4B). For example, the highly abundant transcripts NEAT1 and MALAT1 showed half the protein occupancy after 30 minutes of treatment (adj. p=7.01E-16 and p= 1.60E-05, Figure 4B). Notably, this reduction occurred across the entire body of the transcripts (Figure 6A & Figure S6B). Both MALAT1 and NEAT1 have been reported to bind sites of active transcription (36) suggesting that arsenite led to release of their chromatin interactions during the transcriptional transition from growth and proliferation towards stress and survival. The strong read coverage of MALAT1 for both the pull-down and the input allowed us to use T-C transitions to test for differential binding within the transcript on a nucleotide level. We found two sites in close proximity with particularly decreased protein occupancy for both the 15 and 30 minute time point (>6.0 fold decrease over untreated, adj. p<1.0E-3, Figure 6B). Reportedly, MALAT1 associates in this region with serine/arginine (SR) proteins to modulate alternative splicing by acting as a molecular sponge for hyperphosphorylated splicing factors (37). Since many arsenite response genes such as HSPA1A, HSPA1B, DNAJB1, EGR1, JUN or FOS etc. (Figure 4A) have no or very few introns, this might reflect changes in the RNA splicing behaviour of stressed cells, where overall fewer splicing operations occur. Only a small number of non-coding RNAs increased their protein occupancy during arsenite stress, including the uncharacterized transcript AC019322.4 (2.3 fold increase over untreated, adj. p=0.02, Figure 4B). AC019322.4 gained pull-down read density especially in one region close to its 3’ end, whereas the total expression in the input remained constant (Figure S6B). The gene coding for AC019322.4 resides immediately upstream of an annotated enhancer region and is surrounded by twelve RNA polymerase III genes, including six intact Y and snoRNAs as well as six U6 and 7S pseudogenes in a 500 kbp radius (Figure S6D). For comparison, MALAT1 and NEAT1 have two RNA polymerase III genes in this radius. Transcription of 7S pseudogenes multiplied after 30 minutes of arsenite stress (8.8 fold increase over untreated, adj. p= 5.2E-09, Figure 4A) leading us to speculate that AC019322.4 might promote RNA polymerase III transcription as an enhancer RNA via protein interactions in its 3’ end.

**Figure 6:**
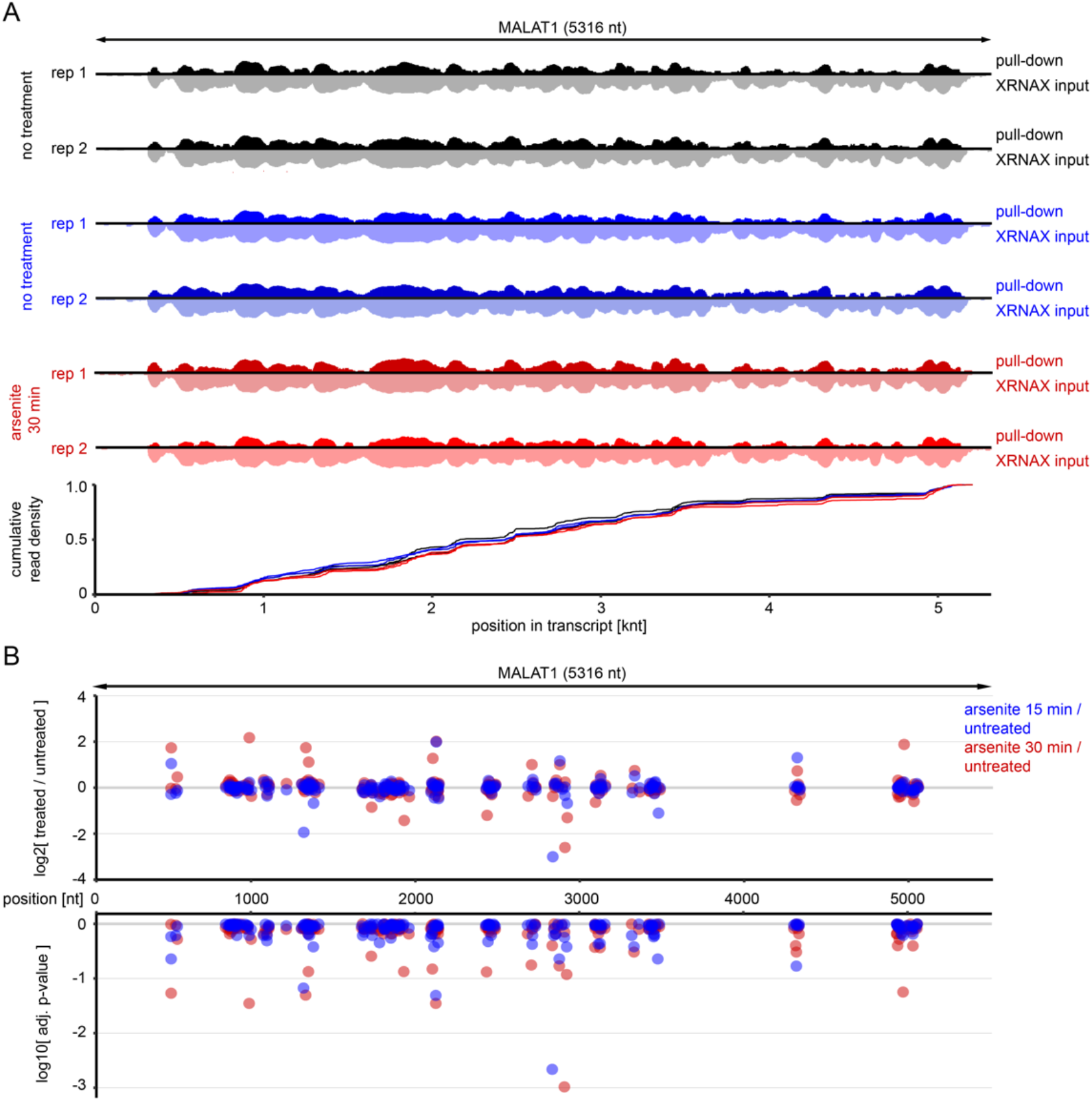
Changes in protein occupancy across the lncRNA MALAT1 during arsenite stress. A) Gene browser view of MALAT1. For each replicate the pair of pull-down (up, dark) and the matched input (down, light) are shown. For better visibility scaling is logarithmic and ranges in all plots from 0-3000 reads. The density plot at the bottom shows the distribution of reads along the transcript for each replicate of the pull-down. B) DEseq results along the sequence of MALAT1. Scatter plot comparing changes in normalized T-C transition frequencies (top) to their significance level (bottom).

In summary, these examples illustrate how PEPseq can be applied to monitor protein interactions with non-coding RNA across many thousands down to a single nucleotide.

## Discussion

### A straightforward RNA sequencing strategy for transcriptome-wide mapping of protein occupancy

Powerful methods exist that interrogate the interactions between a single, known protein towards the transcriptome in order to elucidate hypotheses about a particular post-transcriptional mechanism.(5) However, an unbiased approach such as PEPseq can help to generate new hypotheses about potential post-transcriptional regulation, without requiring prior knowledge about the identity of the proteins involved. An analogy can be drawn to the study of transcription regulation and widely used methods such as DNase-seq or ATAC-seq, that probe chromatin accessibility to identify regulated sites in the genome while staying agnostic to the identity of the proteins that access them (38). Because RNA in contrast to DNA is not covered in histones, PEPseq uses changes in protein-RNA interactions to locate sites of potential regulation. Conceptually, PEPseq is related to protein occupancy profiling (12), however, it introduces a number of key improvements. First, PEPseq uses XRNAX instead of poly-dT enrichment and is therefore able to obtain the protein occupancy of the entire transcriptome and not only the polyadenylated part.(14) Next, PEPseq was designed for straightforward comparisons between conditions and uses an input-controlled quantification implemented with the easy-to-use RNA sequencing tool DESeq2 (23), making it accessible to less-specialized users. We demonstrate that PEPseq provides two quantitative measures, T-C transitions and read counts, which can be applied depending on the use case. While read counts are more sensitive and allow to screen for changes in protein occupancy across the entire transcriptome, T-C transitions provide quasi nucleotide resolution and the ability to pinpoint interaction changes on abundant transcripts. In comparison to the recently published RNP-MaP, which requires cells to be reacted 15 minutes with the NHS-activated crosslinking probe at room temperature followed by ten minutes of UV crosslinking (13), PEPseq requires approximately one minute of UV crosslinking on ice. Consequently, PEPseq has a resolution of minutes and the ability to quantify changes between time points, whereas RNP-MaP has so far been used to probe static interactions within one time point. Additionally, PEPseq starts with XRNAX and therefore offers the potential to investigate protein-RNA interactions both in a proteome-centric and transcriptome-centric manner from the same sample. We imagine future applications of PEPseq in the discovery of post-transcriptional regulatory processes during other perturbations such as the exposure of cells to drugs, infections, or during development.

### PEPseq opens new perspectives on protein-RNA interactions to help hypothesis forming

Using PEPseq we observed immediately discernible differences in protein occupancy between the three functional mRNA regions (i.e. UTRs and CDS, Figure 2). Moreover, changes in protein occupancy across the CDS of mRNA and in the rRNA of the ribosomal peptide exit tunnel were consistent with translational arrest and ribosome stalling in MCF7 cells, what aligned with a shift in read lengths within published Ribo-seq data in HEK293 cells. Thus, our PEPseq analysis pointed out a dominant effect in the transcriptome, which we could confirm in a different cell line using orthogonal methodology. Conceptually, this illustrates the enabling power of PEPseq to generate hypotheses from a global view on protein occupancy across the transcriptome during a particular treatment.

Arsenite exposure has long been known to coincide with activation of the integrated stress response (ISR) and phosphorylation of EIF2α, leading to translational arrest at the stage of translation initiation (31, 32). So far it was assumed that arsenite triggers translational arrest by causing covalent modifications to cysteines and consequent protein misfolding (17), based on toxicological studies where a cysteine-containing methylase detoxifies arsenite to dimethylarsinic acid (39, 40). However, to our knowledge, widespread cysteine modification of the proteome through arsenite has never been demonstrated. In fact, using the mass-tolerant search engine MSfragger (41) we failed to detect an accumulation of any modification in proteomic data from arsenite-treated cells in comparison to untreated cells (data not shown), both in the proteomic data presented in this study as well as in our previous work (14). Moreover, studies in mouse (15) and human cells (42) showed that arsenite does not activate the kinase PERK, which is characteristically triggered by endoplasmic reticulum (ER) stress and the unfolded protein response (43). Therefore, at present it is unclear what exactly leads to activation of the ISR kinases upon arsenite stress. Our findings indicated that the cue might be stalling ribosomes, which, like arsenite, have been reported to be a potent activator of the ISR kinase GCN2 (44). Future studies may investigate if arsenite itself traps 80S ribosomes in a post-translocation conformation, and if in turn they activate GCN2.

### Translational control after stress follows a restart sequence that represses growth and fosters regulation

The biological purpose for the formation of stress granules remains enigmatic, and it is still debated what determines their mRNA composition (29). We show that mRNAs with strong stress granule localization get an immediate kick-start in translation upon stress withdrawal, whereas mRNAs excluded from stress granules remain repressed for some time. Among the included or excluded transcripts, respectively, we identified groups of mRNAs coding for proteins with shared biological functions. This suggests that the sorting of mRNAs into stress granules prepares a biological program, which is put into action when stress signals subsides. We present evidence that during arsenite stress many of these mRNAs collect stalled ribosomes in their CDS, which previously was shown to repel transcripts from stress granules (19, 45). In fact, one defining feature of stress granules is that they aggregate mRNA with stalled pre-initiation complexes and for the most part do not contain 60S subunits (46–48). Thus, we propose a model for the recruitment of RNA into stress granules where on the one hand healthy mRNA without stalled ribosomes is collected, primed to engage in translation as soon as the EIF2α translation initiation block is removed. On the other hand, mRNA carrying stalled ribosomes are excluded from stress granules, and the ribosome blockage in their CDS delays their expression during the initial phase of recovery. We show that after stress this leads to increased expression of DNA-binding and heat shock proteins, whereas expression of ribosomal proteins remains repressed for some time. This intuitively makes sense because after an emergency event has led to translational shutdown, it seems in the best interest of the cell to take control of the stress situation first before investing resources into the expansion of its translational machinery again. Khong *et al*. demonstrated that most of the mRNAs present in the total transcriptome seem to be present in stress granules, although in strongly skewed abundances (Figure S5B) (30). This indicates that the process of stress granule aggregation samples RNA from the entire cytosolic transcriptome, and that sampling does not occur at random as demonstrated by mRNAs coding for cytosolic ribosomal proteins. We speculate that sequence biases such as high GC3 content might serve as evolutionary evolved trip wires for translating ribosomes that lead to increased ribosome stalling on particular sets of transcripts and their exclusion from stress granules. Overall, we propose that stress granules collect healthy mRNA without stalled ribosomes that organize the recovery of the cell when stress is overcome, and when conditions for normal translation are restored.

Our findings illustrate how monitoring changes in protein occupancy with PEPseq can highlight previously unknown protein-RNA interactions to form a working hypothesis for the post-transcriptional response towards a particular treatment. In our case, the increase in protein occupancy across the CDS suggested that this regulation involved ribosomes. In other cases the intersection of PEPseq with eCLIP data (9) could be used to identify candidate proteins potentially responsible for changes in protein occupancy at certain locations of the transcriptome. Thus, PEPseq offers an unbiased discovery platform for the investigation of post-transcriptional processes in differentially treated cells.

## Acknowledgments

We thank Vladimir Benes, Bettina Hase and Nayara Azevedo (Gene-Core EMBL Heidelberg) for RNA sequencing as well as advice and discussion. We thank Charles Girardot (Genome Biology Computational Support) for support with handling and storage of sequencing data. We thank Kazuya Ichihara for additional information about their Ribo-seq dataset.

## Author contributions

Conceptualization: JT, JK; Methodology: JT; Investigation: MJ, EB, JT; Visualization: JT, MJ; Funding acquisition: JK; Administration: JT, JK; Supervision: JK, CD; Writing – original draft: JT, JK; Writing – review & editing: JT, EB, MJ, JK, CD

All authors declare that they have no competing interests.

## Data availability

Sequence data from PEPseq libraries generated in this study have been submitted to EMBL-EBI ENA under the accession PRJEB58258. Proteomics data is currently submitted to PRIDE.

**Figure S1:**
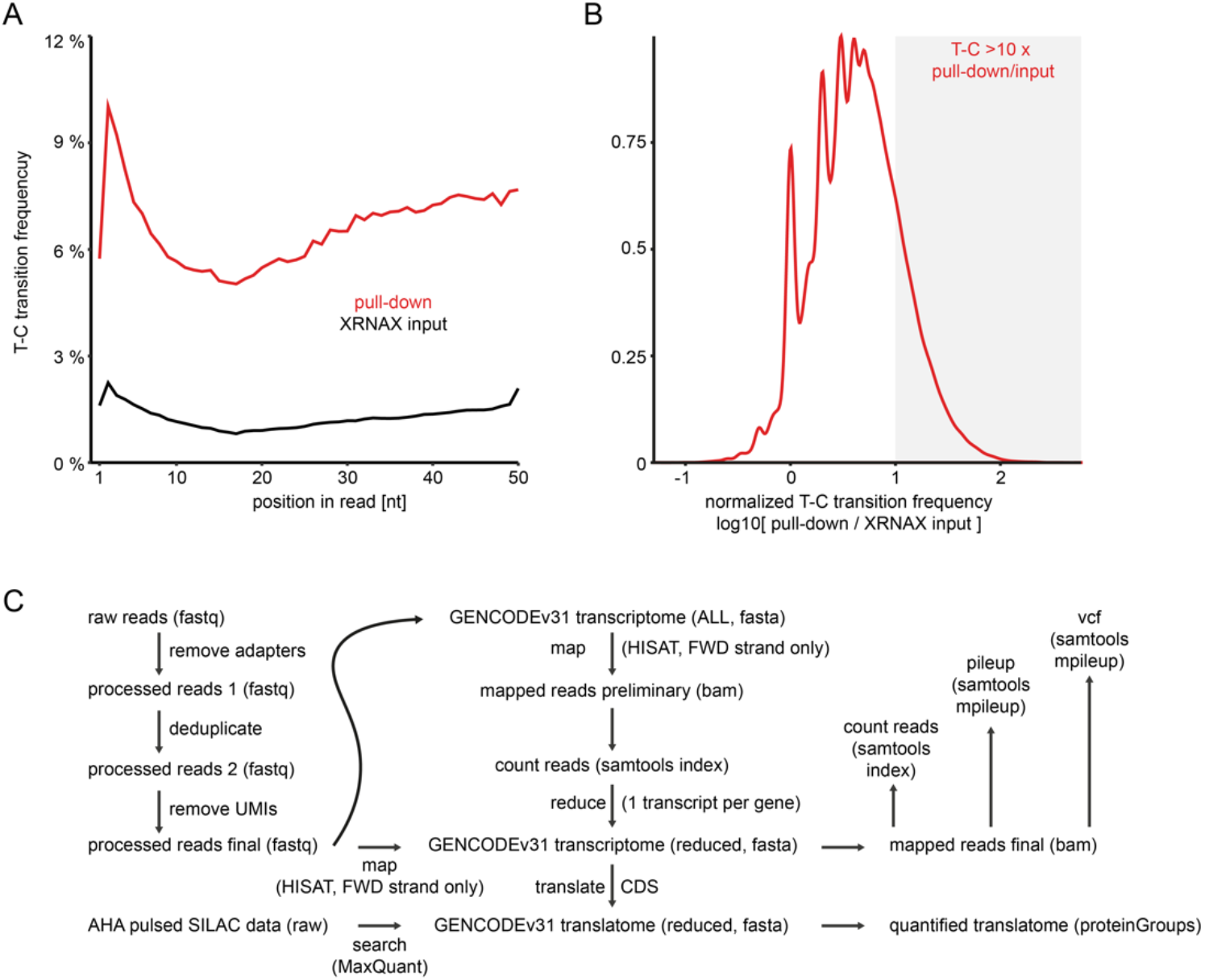
Properties of PEPseq RNA sequencing reads and mapping strategy. A) Line graph comparing T-C transition frequencies within reads from peptide-enriched fragments (pull-down) and the matched input they were extracted from (XRNAX input). B) Density plot showing ratios of T-C transition frequencies between pull-down and XRNAX input (normalized T-C transition frequency). T-C transition frequencies were determined by counting T-C transition events at each T position in the transcriptome and comparing these to the read coverage at the same position. C) Processing pipeline for the PEPseq libraries and MS-based proteomics. Left: Raw reads are freed from barcodes and deduplicated in order to remove PCR duplicates. Subsequently, UMIs are removed from both ends to yield the final processed reads used for quantification and T-C calling. Middle: A custom transcriptome is generated from the GENCODE v31 transcriptome, with the goal to have each GENCODE gene represented by a single transcript, coding for a single protein isoform. Therefore, reads from all libraries (pull-down and XRNAX input, all replicates from all timepoint) are combined and mapped against the initial GENCODE v31 transcriptome containing all transcriptome isoforms (ALL). For all isoforms reads are counted in order to identify the one transcript with the most mapped reads. In case of identical read counts, the shortest isoform is used. Right: After the optimal transcriptome has been determined, containing one transcript isoform representing each GENCODE gene (GENCODEv31 transcriptome (reduced)), the individual libraries are mapped. Subsequently, alignments are used to generate read counts per transcript, pileup and vcf files using samtools. Protein coding transcripts contained in the reduced transcriptome are translated to proteins according to the annotation of their CDS and used to search proteomics data.

**Figure S2:**
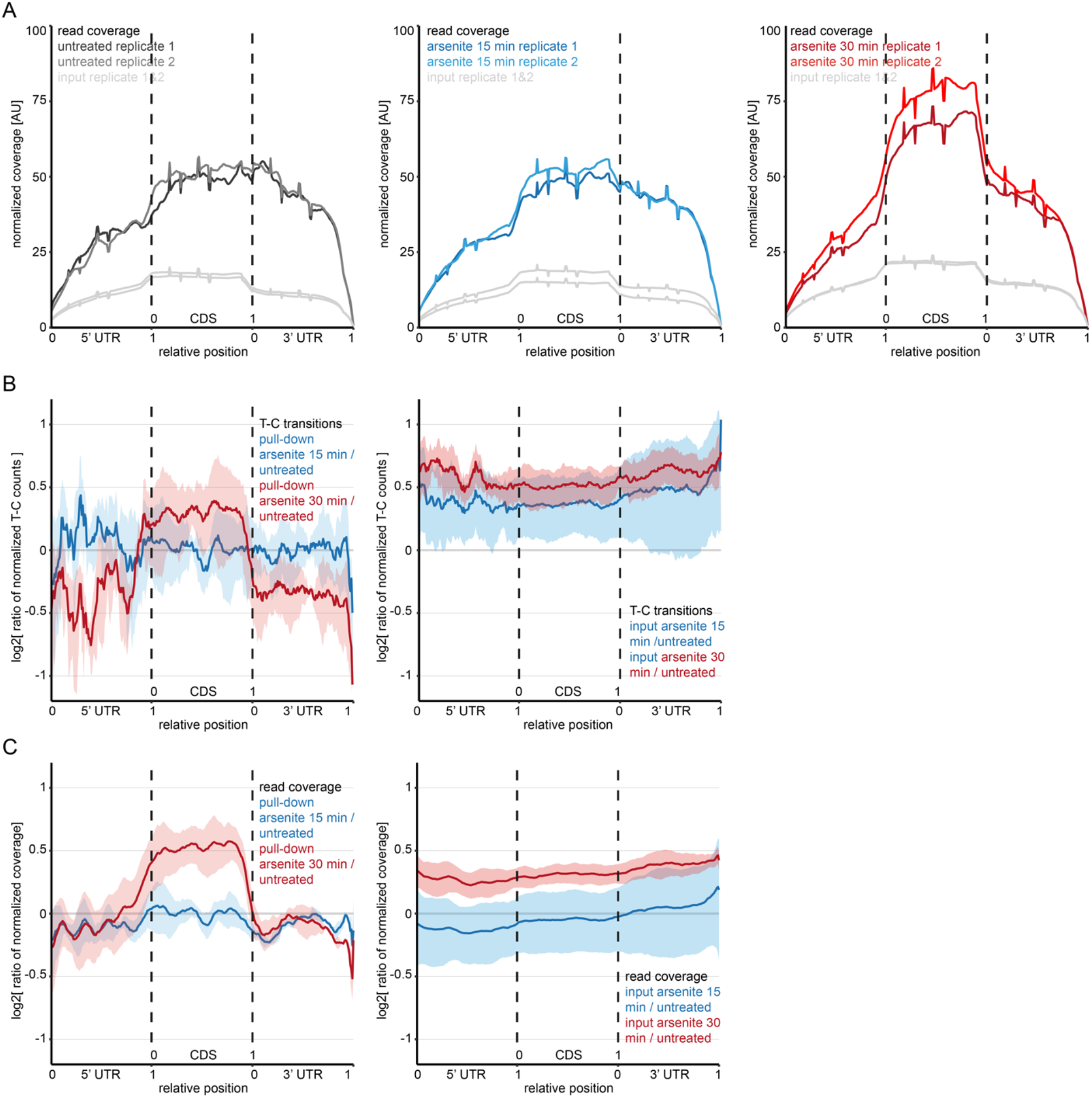
Changes in protein occupancy across mRNA upon arsenite-induced translational arrest. A) Metagene plot as in Figure 2A but for read coverage. Normalization occurred in the same way as for T-C transitions. B) Metagene plots illustrating the change in T-C transitions upon arsenite stress across all detected mRNAs. Thick lines indicate the ratio means compared to untreated MCF7 cells, shaded areas indicate one composite standard deviation. Normalization occurred as in A. Compared are the pull-down (left) to the XRNAX input (right), which are combined into one graph in Figure 2B. C) Same as in B but for read coverage instead of T-C transitions. Compared are the pull-down (left) to the XRNAX input (right), which are combined into one graph in Figure 2C.

**Figure S3:**
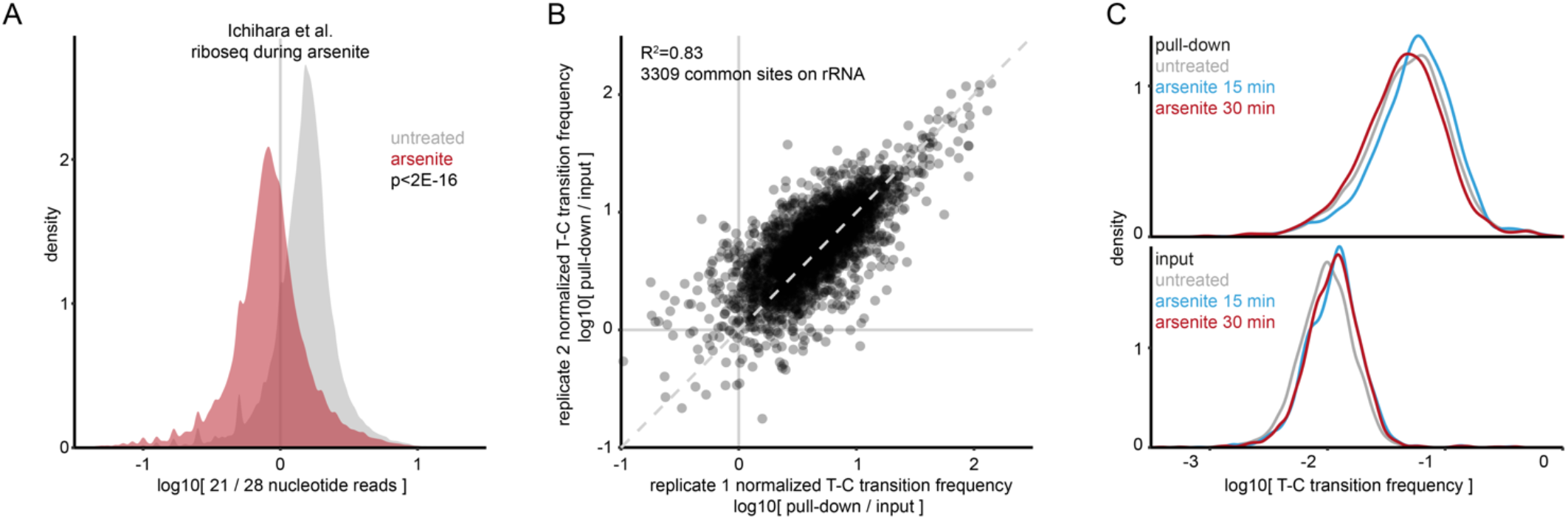
Effects of arsenite stress on Ribo-seq and PEPseq data. A) Density plot illustrating the ratio of 21 nucleotide to 28 nucleotide ribosome footprints per transcript in published Ribo-seq data (Ichihara *et al*.) from untreated (grey) to 60 minutes arsenite-treated U2OS cells (red). B) Scatterplot comparing normalized T-C transition frequencies on rRNA between replicates. Each point represents the normalized T-C transition frequency at one particular site and time point compared between duplicates. R^2^ indicates the Pearson correlation. B) Density plot comparing T-C transition frequencies on rRNA between time points. C) Density plots comparing T-C transition frequencies between time points for the pull-down (top) and the input (bottom).

**Figure S4:**
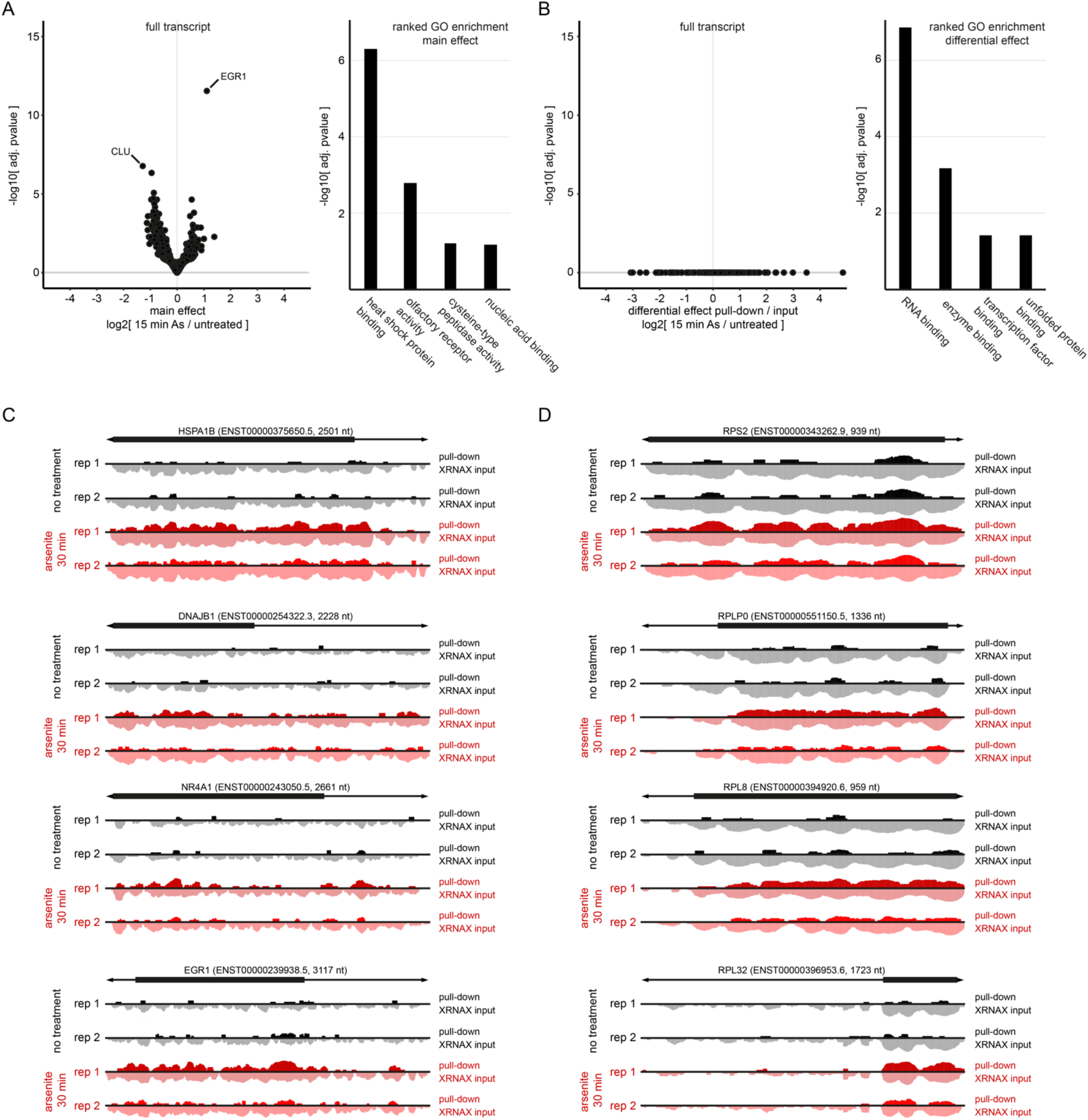
Arsenite stress leads to distinctive changes in protein occupancy across specific transcripts. A) & B) Same analysis as in Figure 4A & B but using the 15-minute timepoint. C) Gene browser views of four exemplary transcripts identified in Figure 4A. For each replicate the pair of pull-down (up, dark) and the matched input (down, light) are shown. The bold area of the upper arrow indicates the CDS of each transcript. For better visibility scaling is logarithmic and ranges in all plots from 0-400 reads. D) Same as in C but for exemplary transcripts coding for cytosolic ribosomal proteins identified in Figure 4C.

**Figure S5:**
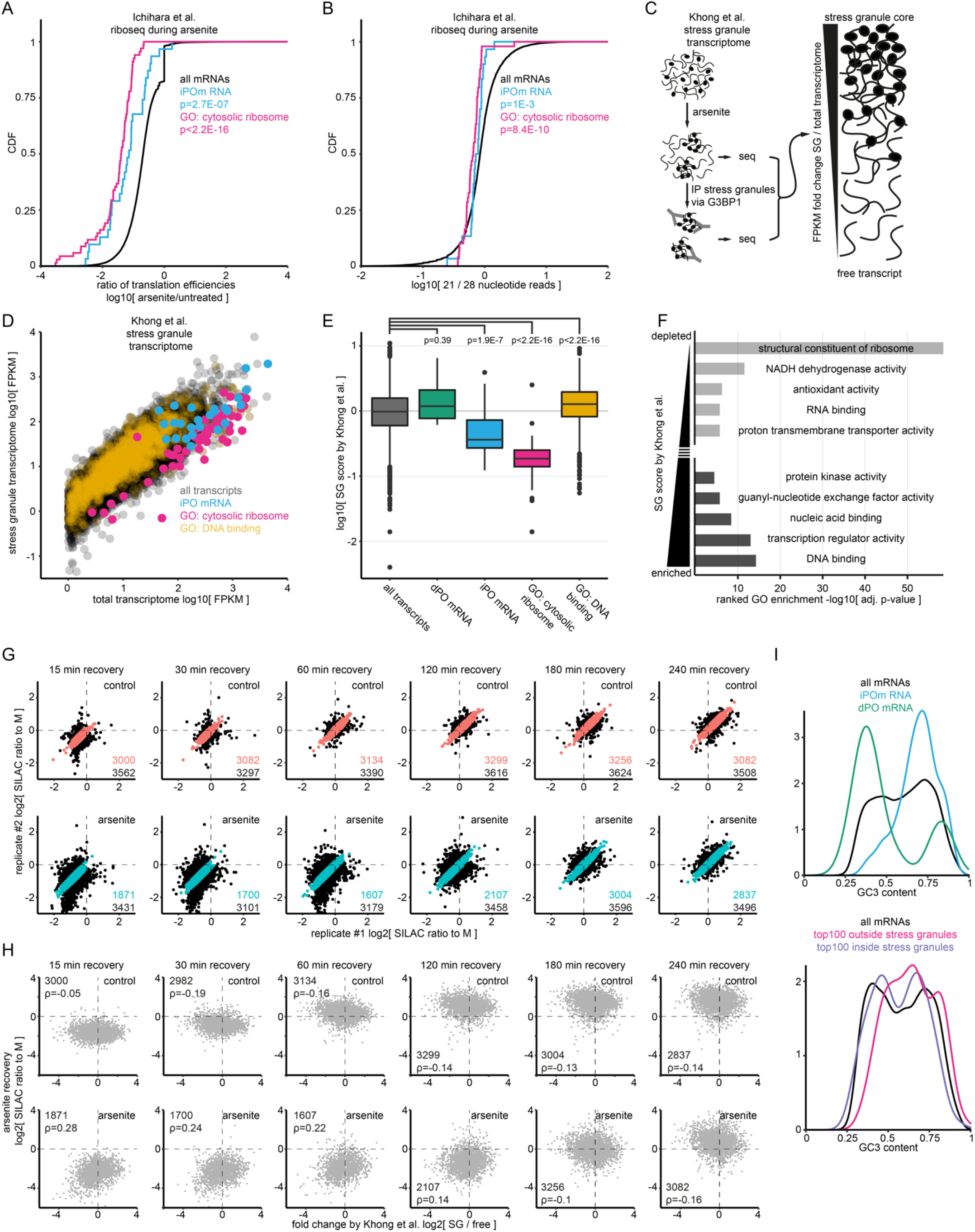
Protein production during and after arsenite-induced translational arrest. A) Cumulative distribution of relative translation efficiencies reported by Ichihara *et al*. using Ribo-seq. Untreated HEK293 cells are compared to cells treated with arsenite for one hour. Arsenite concentrations were 10 times lower (40 μM) than in this study (400 μM). P-values indicate results for a two-sided Kolmogorov-Smirnoff test between the indicated group of mRNAs and all mRNAs. B) Cumulative distribution for the ratio of 21 nucleotide to 28 nucleotide ribosome footprints per transcript from the same data as in A but only for arsenite treated cells. Testing occurred as in A. C) Experimental scheme followed by Khong *et al*. to derive stress granule fold changes. U2OS cells expressing GFP-tagged G3BP1 were subjected to arsenite to induce stress granules. Subsequently total RNA was extracted for RNA sequencing of the total transcriptome and stress granules purified via immunoprecipitation for RNA sequencing of the stress granules transcriptome. D) Scatter plot illustrating published transcript abundances by Khong *et al*. from the total transcriptome and the stress granule transcriptome of U2OS cells. Each point represents one transcript/gene. E) Boxplot comparing the enrichment of transcripts in stress granules reported for 11195 genes by Khong *et al*.. Higher foldchanges indicate stronger localization to stress granules as observed in U2OS cells. Testing occurred between all reported genes (all), iPO mRNAs and subgroups with the indicated gene ontology annotation using a two-sided Kolmogorov-Smirnov test. F) Ranked gene ontology enrichment analysis for proteins inside (top, dark grey) and outside (bottom light grey) of stress granules. For each analysis genes were sorted in descending (top) or ascending (bottom) order of the fold change reported by Khong *et al*.. Non-redundant, representative terms were curated to capture the top hits of each analysis. G) Scatter plots comparing biological replicates with SILAC label swap for quantification of nascent protein during the recovery from arsenite. Untreated control cells (top, red) are compared to cells treated with arsenite for 30 minutes (bottom, cyan). Each dot represents one protein, coloured proteins lie within REM<30% between replicates. Black number indicates proteins detected in both replicates, coloured number within REM<30%. H) Scatter plots comparing the stress granule score by Khong *et al*. to the relative protein production determined by AHA-labelling during recovery. Relative protein production rates are means of duplicates displayed in A filtered for REM<30%. Number indicates the compared points and ρ the Spearman correlation. I) Density plots comparing the GC3 content of mRNA groups. For visibility transcripts discovered by PEPseq (top) and transcripts with extreme stress granule localizations according to Khong *et al*. (bottom) are displayed in different panels. GC3 content was calculated for all mRNAs in our reduced GENCODEv31 transcriptome containing one mRNA per gene (see Figure S1C).

**Figure S6:**
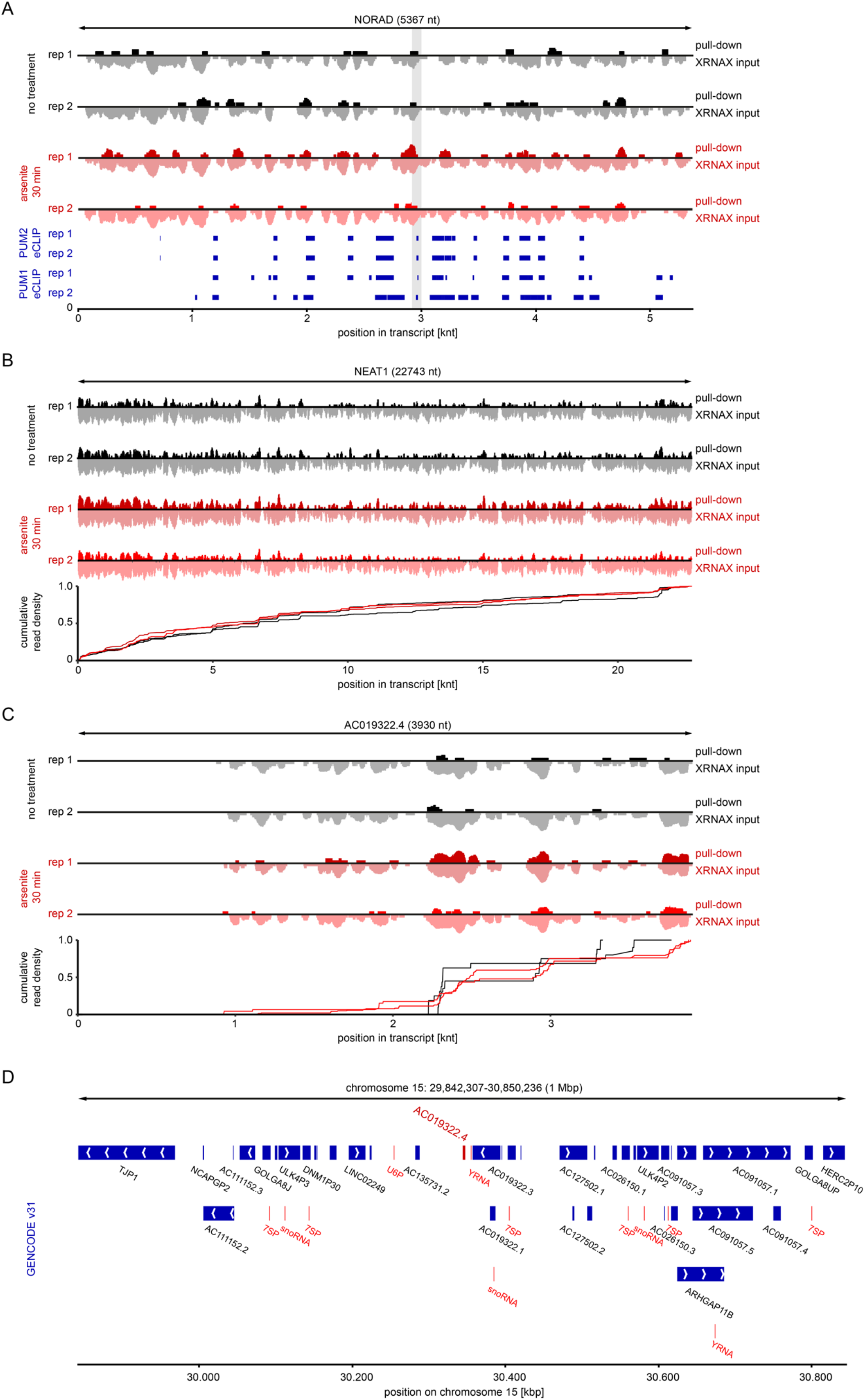
Changes in protein occupancy across non-coding RNA upon arsenite stress. A) Gene browser view of NORAD. For each replicate the pair of pull-down (up, dark) and the matched input (down, light) are shown. For better visibility scaling is logarithmic and ranges in all plots from 0-70 reads. In addition published eCLIP data from K562 cells for the PUM1 and PUM2 are displayed and overlapping changes in read density highlighted in grey. B) Gene browser view of NEAT1. For each replicate the pair of pull-down (up, dark) and the matched input (down, light) are shown. Same as in A but scaling ranges from 0-500 reads. The density plot at the bottom shows the distribution of reads along the transcript for each replicate of the pull-down. C) Gene browser view of AC019322.4. Same as in A but scaling ranges from 0-120 reads. D) Gene browser view for the AC019322.4 locus on chromosome 15. Displayed is a 500 kpb window upstream and downstream of the AC019322.4 gene (dark red) and the GENCODE v31 annotation. RNA polymerase III transcripts are highlighted in light red.

